# Neuronal targeted AAV Micro-Dystrophin restores neurobehavioural co-morbidities, grip strength and motor coordination in *mdx52* mouse model of Duchenne Muscular Dystrophy

**DOI:** 10.64898/2026.05.22.727055

**Authors:** Konstantina Tetorou, Monica Rebeca Gil Garzon, Darren Chambers, M Görkem Özyurt, Filipe Nascimento, Wing Sum Chu, Simon N. Waddington, Benjamin W. Jarvis, Alex Kavanagh, Nicha Songsliph, Joanne Ng, Francesco Muntoni

## Abstract

Duchenne muscular dystrophy (DMD) is a X-linked disorder caused by mutations in the *DMD* gene, which disrupts production of multiple isoforms of dystrophin in multiple organs namely muscle, heart and brain. While progressive muscle disease and cardiomyopathy are the hallmarks of DMD, over 40% of individuals also experience significant neurobehavioral comorbidities, including autism spectrum disorder (ASD), attention deficit hyperactivity disorder (ADHD), obsessive compulsive disorder (OCD) and intellectual disability. These deficits are linked to the loss of brain isoforms and approximately 90% of DMD individuals have loss of either Dp427 or both Dp427 and Dp140 in the brain.

We studied the *mdx52* mouse model, which lacks these isoforms and exhibits severe fear and anxiety-like behaviours, suitable to evaluate the therapeutic efficacy on neuro-comorbidities after delivering a neuronal-targeted adeno-associated virus (AAV) micro-dystrophin (µDys) therapy. We compared two delivery routes intravenous (IV) and intracerebroventricular (ICV) in neonatal *mdx52* male mice to assess impact on an extensive range of neurobehavioural aspects including emotional reactivity, neurocognitive, OCD and motor coordination deficits. While both routes successfully reduced emotional reactivity and anxiety-related behaviours, IV delivery emerged as the superior therapeutic strategy addressing a more comprehensive spectrum of DMD related brain co-morbidities.

Critically, significant improvements in cognitive deficits and OCD-like behaviours were achieved only through IV delivery. This was associated with a widespread lower transduction pattern across the brain, including hindbrain and cerebellum, which were less effectively targeted by ICV injection, although forebrain transduction with ICV delivery was higher.

Brain µDys expression successfully restored dystrophin interactors dystroglycan, syntrophin and pre- and post-synaptic functional interactors VGLUT1, gephyrin, GABA_A_R with both delivery methods. These results demonstrate that while ICV gene therapy results in improved emotional reactivity and anxiety-related behaviour in the *mdx52*, only the systemic, neuronal-targeted gene therapy efficiently transduced the central nervous system restoring neuronal synaptic functional complexes of both Dp427 and 140 isoforms and simultaneously restored peripheral NMJ dystrophin deficiency.

Beyond cognitive restoration, while both routes improved aspects of gait on CatWalk XT, only IV delivery significantly enhanced motor coordination on the Beam walk and, unexpectedly, normalised grip strength. This was specifically linked to the selective expression of µDys at neuromuscular junctions (NMJs), which corrected post-synaptic electrophysiological dysfunction of *mdx52* mice. Our findings establish a significant foundation for incorporating brain-directed strategies into the future therapeutic approaches for DMD, offering a holistic approach to treating DMD as a multisystemic disease.

## Introduction

Duchenne muscular dystrophy (DMD) is a severe, progressive, X-linked recessive neuromuscular disorder primarily affecting boys with a prevalence of 1 in 3500 to 1 in 5000 live male births^1^, and global incidence of approximately 3.6 per 100,000 people.^2^ The *DMD* gene spans 2.3 Mb and consists of 79 exons with 7 internal promoters,^3^ each linked to unique first exons giving rise to at least 7 proteins of different sizes that are expressed or co-expressed in different tissues or organs (**Fig 1A**). Pathogenic mutations in the *DMD* gene result in the absence of Dystrophin protein that protects muscle from progressive damage.^4^ The full-length dystrophin protein (427 kDa) is expressed under the sarcolemma in cardiac and skeletal muscle cells, with a crucial role in linking the extracellular matrix to the cytoskeletal actin via a transmembrane complex associated to dystroglycan (dystrophin associated protein complex DAPC), providing resistance to mechanical shear forces during muscle contraction and relaxation.^5^

**Figure 1.**
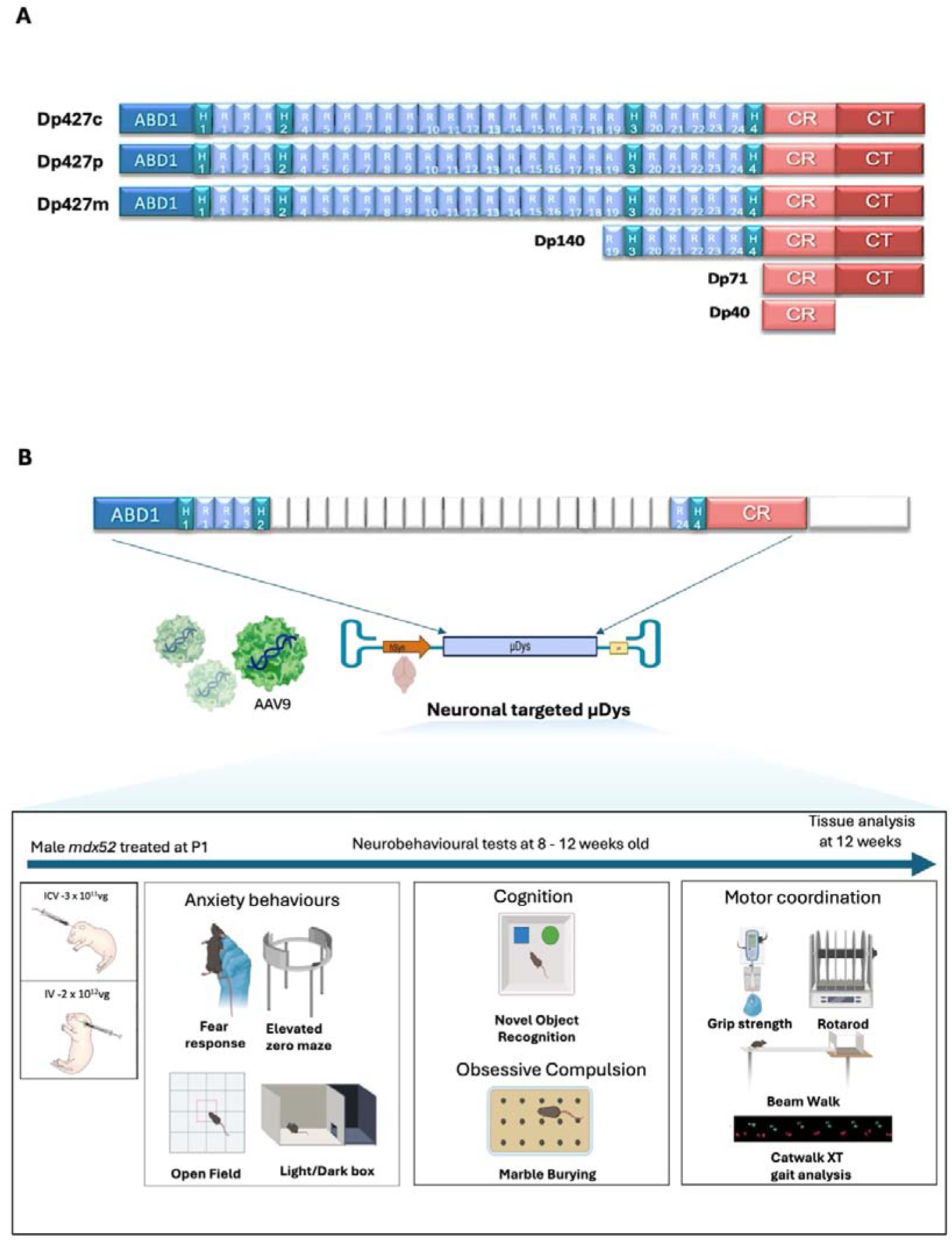
Illustration of gene therapy for brain dystrophin deficiency. **(A)** Schematic of dystrophin isoforms expressed in the brain Dp427c cortex, Dp427p Purkinje, Dp427m muscle, Dp140,71 and 40. (**B)** Experimental plan: *mdx52* P1 mice were injected ICV and IV with AAV9.hSyn.µDys and were subjected to a battery of behavioural tests along with control WT and *mdx52* untreated (n=14-15 per group). The behavioural tests included fear response, elevated zero maze, open field and light/dark box measures of anxiety, novel object recognition test as a measure of cognitive ability and recognition memory, marble burying as a measure of anxiety and OCD behaviours, and rotarod, grip strength and Beam Walk Test, CatWalk XT assess motor coordination and muscle function. The tests were performed between 8-12 weeks post-treatment allowing 24-48 hour intervals between assessments.

DMD neuromuscular and cardiac manifestations lead to rapid progression of weakness, loss of ambulation and cardiorespiratory insufficiency resulting in shortened lifespan of approximately 30 years.^6–8^ Early reports indicated that 30% of DMD children have intellectual disability^9^ and more recent studies have demonstrated that neurobehavioral co-morbidities are also frequently found in DMD, including attention-deficit/hyperactivity disorder (ADHD), autism spectrum disorder (ASD), emotional disorders, obsessive-compulsive disorder (OCD),^10–12^ anxiety and increased unconditioned startle response.^13,14^ These co-morbidities cumulatively affect more than 40% of the DMD population, are associated with markedly reduced independence, quality of life, and impact on prognosis and survival.^15,16^ Knowledge generated also by our group has clarified that the variable clinical brain involvement is strongly related to the location of the *DMD* mutation and its consequences on the dystrophin isoforms expressed.^17,18^

Multiple dystrophin isoforms are expressed in the brain: in addition to the full length Dp427, also expressed in muscle, they include Dp140, Dp71, each driven by independent promoters and with different expression profiles in human and mouse brain.^17,19–22^ These isoforms are related to neuronal development, synaptic transmission, connectivity and blood brain barrier integrity, through a diversity of protein complexes.^23^ To add complexity, the full length Dp427 isoform is driven by 3 independent promoters, each linked to a unique short first exon: Dp427c (cortical), Dp427p (cerebellar Purkinje) in addition to the Dp427m (muscle) isoform.

All DMD individuals have complete loss of the 3 full length Dp427 isoforms in both muscle and brain, independent to *DMD* mutation location. Mutations located between exons 51 and 62 also abolish the production of Dp140, while those downstream to exon 63 also affect Dp71, hence resulting in complete loss of each of the DMD isoforms (**Fig. 1A**). Dp140 promoter is located in intron 44, while exons 45-50 are part of its 5’UTR. This suggests that mutations located between exons 45 and 50 should also affect Dp140 production, but as this isoform is exclusively expressed in the brain, formal demonstration of the extent to which these mutations affect Dp140 production in patients is not known. These shorter isoforms show different spatiotemporal expression, related to brain development and are expressed in different brain cellular sub-types.^17,22^

Brain Dp427 is involved in the localisation and functioning of postsynaptic γ-Aminobutyric acid type A (GABA_A_) receptors in mouse, with clear defects observed in DMD mouse models.^24^ Dp140 shows higher expression during fetal stages and lower post-natal levels.^17,19,20,22^ Recent studies have indicated that the Dp140 loss is associated with abnormalities in glutamatergic neurotransmission in the *mdx52* mouse model.^25,26^ This model is deficient in both Dp427 and Dp140, but has intact Dp71.

The Dp71 isoform is the most abundantly expressed isoform in the human and mouse brain with temporal and cell-type specific expression during both development and in adult stages.^17,19,20^ This isoform is mainly associated with aquaporin in the brain and regulation of ion exchange but also contributes to glutamatergic synaptic organisation and function.^27^

The cumulative deficiency of different brain isoforms secondary to the location of the causative *DMD* mutation, has provided us with critical insight for their contribution to brain function: the isolated deficiency of Dp427 (∼ 40% of cases) does not significantly affect intellectual function but is associated with increased risk of neurobehavioural comorbidities; the combined deficiency of Dp427 and Dp140 (∼50-55% of cases) leads to worsening of the neurobehavioural complications and is associated with a mean IQ of 75.^28^ The deficiency of all brain isoforms observed in 6-10% of patients is associated with the most severe neurocomorbidities and a mean IQ of 55.^29–34^

Despite the increasing evidence linking dystrophin deficiency to neurocognitive and neurobehavioural deficits, there have been few approaches to address brain dystrophin deficiency relative to the genetic therapies developed for dystrophin restoration in muscle. These genetic therapies include FDA approved antisense oligonucleotides (ASO) and adeno-associated virus (AAV) micro-dystrophin gene therapy and multiple other genetic therapies in clinical trials.

ASOs produce an internally deleted but in-frame mRNA by removing exons adjacent to out-of-frame deletions, hence restoring the *DMD* reading frame and resulting in expression of a shortened but functional dystrophin. Four ASO drugs targeting different DMD mutations have received FDA conditional approval (eteplirsen, golodirsen, casimersen and viltolarsen, targeting exon 51, 53, 45 and 53 respectively). However, these ASOs are administered intravenously and do not cross the blood brain barrier, and do not address brain dystrophin deficiency.^35,36^

Several studies have used direct delivery of ASO to restore dystrophin production in the brain of the *mdx* mouse achieved partial phenotypic improvement.^37,38^ However inefficient biodistribution irrespective of the different direct brain administration routes and dose-related toxicity limit their potential for clinical translation.^39,40^

An alternative approach is gene supplementation using AAV vectors to deliver a therapeutic transgene to restore dystrophin function.^41^ The large size of *DMD* mRNA (11.5kb) is beyond AAV packaging capacity (maximum 4.7kb).^42,43^ Significant effort has been devoted in the last 2 decades in engineering shortened but functional micro-dystrophins (µDys), containing ∼1/3 of the gene, but retaining essential domains for muscle function.^44^ All µDys lack the majority of the 24 spectrin like repeats, contain two or three of the four Hinge domains, and have only a portion of the C-terminus.^45,46^ All AAV gene therapies under development for DMD, including recently FDA approved AAVrh74 ELEVIDYS (delandistrogene moxeparvovec-rokl) are delivered intravenously, do not cross the blood brain barrier and use muscle specific promoters thus are not suitable to address brain dystrophin deficiency.

Our study is the first to evaluate the role of neuronal targeted AAV-mediated µDys gene supplementation as a therapeutic approach for brain dystrophin deficiency in an animal model of DMD. By comparing ICV and IV delivery we demonstrate surprising superior efficacy with IV delivery that was able to address a comprehensive spectrum of DMD related brain co-morbidities due to improved widespread transduction. An unexpected effect with IV AAV9.hSyn.μDys was normalisation of grip strength due to transgene expression at the NMJ, where it normalised the NMJ dystrophin defect characteristic of the *mdx* mice resulting in normalised grip strength and beam walk.

While µDys gene supplementation cannot recapitulate the complex pattern of expression of multiple isoforms in the brain, we demonstrate that µDys transgene (similar to the FDA approved ELEVIDYS) (**Fig. 1B**) driven by human synapsin promoter (hSyn) was capable of restoring the known brain dystrophin-associated protein complex (DAPC) for dystrophin brain function through known interacting epitopes expressed in µDys and also identified novel putative interacting domain with syntrophin beta 1. Our findings provide new insights on the potential for incorporating brain-directed strategies into the future therapeutic approaches for DMD.

## Methods

### AAV vector

AAV expression cassette was generated in-house to express μDy*s* transgene (cDNA 3591bp) currently used in FDA approved ELEVIDYS gene therapy for DMD^47^ under transcriptional control of hSyn promoter for neuronal selective expression with downstream polyA. The μDy*s* contains Actin binding domain, H1, R1-3, H4 and Cysteine Rich Domain (CRD) (ΔR4-23/ΔCT) termed μDys (**Fig. 1B**). The expression cassette was packaged into AAV9 by standard triple transfection methods and purified on AKTA GO HPLC systems with POROS Capture Select AAVX as previously described.^48^ Vector was titred by TaqMan^TM^ qPCR to the transgene at 2.7×10^14^vg/mL. An equivalent AAV9.hSyn.GFP vector was produced at 6.3 ×10^13^vg/mL to further delineate biodistribution with neonatal ICV and IV delivery.

### Neonatal intracerebroventricular and intravenous vector injections

All animal procedures were performed in compliance with UK Home Office regulations and the Animal (Scientific Procedures) Act of 1986, and within the guidelines of University College London ethical review committee. The *mdx52* mice colony was mated to generate neonatal mice for study.^49^ Pups were sexed and genotyped at P0 and *mdx52* and male wild-type (WT) littermates were used for experimental studies. AAV9.hSyn.μDys was delivered to neonatal male *mdx52* mice at P1 by intracerebroventricular (ICV) injection at 3×10^11^vg/pup dose, termed *mdx52 ICV*, or delivered intravenously (IV) at 2×10^12^vg/pup via the superficial temporal vein^50,51^, termed *mdx52 IV*. Control male *mdx52* and male WT served as littermate controls (*n=14-15* per group). These mice were monitored for general welfare and underwent behavioural analysis at 8-12 weeks with timed euthanisation 12 weeks post gene therapy for tissue analysis (**Fig. 1B**). AAV9.hSyn.GFP was delivered to wild-type P1 pups for biodistribution study with 33G Hamilton needle and syringe by ICV dose of 5×10^10^ vg/pup, or by IV injection via the superficial temporal vein dose 4[×[10^11^vg/pup (n=4 per group with uninjected littermates as controls). Pups were euthanised for GFP biodistribution analysis at 8 weeks as previously described.^48^ For AAV9.hSyn.GFP biodistribution studies WT dams (Charles River) were time-mated to generate wildtype P1 litters.

### Behavioural assessments

All animals were acclimatised to behavioural room for at least 15 minutes and assessors were blinded to genotype and treatment. The battery of behavioural testing consisted of seven consecutive tests performed with at least a 24-hour interval. To assess effects of neuronal expressed μDys, the 4 groups (*mdx52 ICV n=14, mdx52 IV n=14, controls mdx52 n=15, WT n=15*) were subject to established neurobehavioural assessments for emotional-related and motor deficits observed in *mdx52* mice.^52,53^ These included emotional reactivity (freezing response, light/dark box, open field, elevated zero maze), motor coordination (grip strength, rotarod, beam walk and CatWalk XT), cognitive test novel object recognition (NOR) and OCD with marble burying (**Fig. 1B**).

### Co-immunoprecipitation

μDys interaction with brain interactors was identified using co-immunoprecipitation as previously described^54^ using the MANEX1011B monoclonal antibody that recognises full-length dystrophin and μDys.

### Computational protein-protein complex modelling

Protein structures and protein-protein complexes were predicted using AlphaFold (version 3). Amino acid sequences corresponding to full-length dystrophin and syntrophin ß1 were obtained from UniProt, while the μDys transgene sequence was used as designed in this study included Actin binding domain, H1, R1-3, H4 and Cysteine Rich Domain (CRD) (ΔR4-23/ΔCT). For each prediction, the top-ranked model (ranked_0), based on confidence metrics (pLDDT and PAE), was selected for analysis. Structural visualisation and interface analyses were performed using iCn3D, with interacting residues identified based on spatial proximity.

Schematic interaction networks were generated to summarise residue-level contacts observed at the predicted interfaces.

### Repetitive nerve stimulation (RNS)

Mice were anaesthetised with isoflurane (4% for induction, 2.5% maintenance). The right hindlimb was shaved, ophthalmic ointment applied to prevent corneal drying, and the animal positioned prone within a custom Faraday cage with fore- and hindlimbs extended and separated. Recording electrodes consisted of two multi-stranded perfluoroalkoxy-coated stainless-steel wires (25.4µm; A-M Systems, USA) with fish-hooked exposed tips to anchor them within the muscle belly, inserted into the tibialis anterior (TA) via a 25G needle. For peripheral nerve stimulation, two additional wires were inserted subcutaneously, one proximal to the popliteal fossa and one at mid-thigh, to target the sciatic nerve. The tail was grounded to the Faraday cage.

Sciatic nerve stimulation was delivered with a DS3 constant current stimulator (Digitimer, UK). Bipolar TA EMG signals were amplified (Amplifier and Preamplifier Model MA 103, University of Cologne, Institute of Zoology, Büschges Group, Germany), bandpass filtered between 2Hz and 5kHz and sampled at 20kHz via a CED Power1401 system running Spike2 v8 (Cambridge Electronic Design, UK). Stimulus specificity was verified by visible ankle flexion at suprathreshold intensities together with a compound muscle action potential (CMAP) corresponding to a direct motor response (M-response) in TA at low intensities (∼1–1.5× threshold). 200µs square current pulses were applied to evoke direct responses via orthodromic stimulation, and peak-to-peak CMAP amplitudes were quantified. The largest TA CMAP defined as Mmax, and stimulus intensity was subsequently fixed at 10% above Mmax level to ensure reliable motor axon recruitment. Repetitive nerve stimulation was then delivered at 3, 10, 20 and 40Hz, each as a train of 20 pulses, in a random order for each animal.

### Statistical analyses

Statistical analysis was performed on GraphPad Prism version 10.4.1. The results are expressed as mean ± standard error of the mean (SEM). Data normality was assessed using Anderson-Darling (A2*), D’Agostino-Pearson omnibus (K2), Shapiro-Wilk (W), and Kolmogorov-Smirnov tests. An unpaired two-tailed t-test or one-way ANOVA was used for normally distributed datasets, followed by a post-hoc Dunnet’s or Tukey’s multiple comparison test when comparing means to a control group. In all the analyses, a p-value of less than 0.05 was considered significant.

## Results

### Neuronally targeted AAV9.**μ**Dys ameliorates emotional reactivity in *mdx52* with additional efficacy in cognitive and OCD behaviours achieved only with IV delivery

We firstly applied well-characterised *mdx52* emotional reactivity behaviours as measures of therapeutic responses comparing ICV or IV AAV9.hSyn.μDys administration.

Similar to previous reports, there was no tonic immobility in WT mice following brief scruff-restraint, whilst control *mdx52* showed 43% tonic immobility. Both treatment modalities were efficacious, the ICV treated *mdx52* showed 35% reduction (*P* = 0.0122); with IV treated mice demonstrated superior efficacy with normalisation of the pathological freezing response (9%, *P* = 0.0009) (**Fig. 2A**).

**Figure 2.**
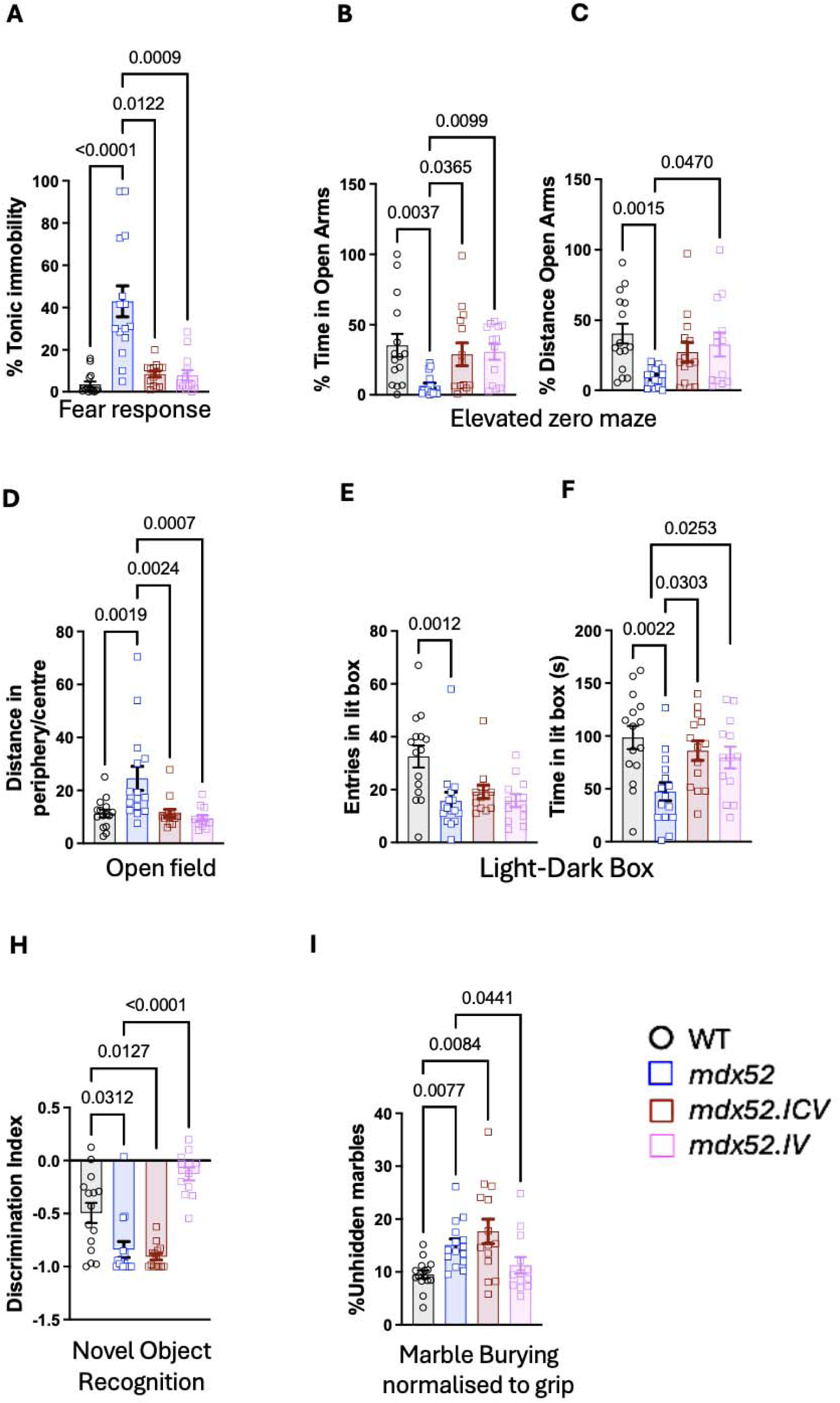
Neuronally targeted AAV9.μDys ameliorates emotional reactivity in *mdx52* with additional efficacy in cognitive and OCD behaviours achieved only with IV delivery. **(A)** Unconditioned fear response as %time spent in tonic immobility **(B)** Elevated zero maze -open arms **(C)** Elevated zero maze -distance **(D)** Central zone to peripheral zone open field distance **(F)** Light-dark box entries to lit compartment **(G)** Light-dark box-time spent inside lit area **(H)** Novel Object Recognition Discrimination index **(I)** % unhidden marbles in marble burying normalised to grip strength. Groups sizes n=14-15. Data are presented as mean ± SEM analysed with Kruskal-Wallis test with Dunn’s post-hoc.

On elevated zero maze, %time spent in open arms was restored to WT levels in all treated mice irrespective of administration method (ICV *P* = 0.0337, IV *P* = 0.0167), while the distance in the open arm was significantly improved only with IV treatment (*P* = 0.0470) (**Fig. 2B, C**).

In open field, proxy for thigmotaxis anxiety behaviour measured as distance in the periphery/central zone was significantly reduced in *mdx52* to WT (*P* = 0.0019). Following gene therapy, % central distance was significantly increased and restored to WT levels following both ICV (*P* = 0.0024) and IV (*P* = 0.0007) delivery (**Fig. 2D, Supplementary Fig 1A**).

In the light/dark box, we measured entries in light compartment and time spent exploring the lit anxiogenic compartment, compared to secure dark compartment. The number of entries in the lit box was significantly reduced in *mdx52* compared to WT (32.5 to 15.6 entries) and with no significant effect of gene therapy irrespective of delivery route. The time spent in lit compartment was significantly reduced in *mdx52* to WT; importantly μDys treatment significantly increased time spent equivalent to WT levels following both ICV (*P* = 0.0302) and IV delivery (*P* = 0.0253) (**Fig. 2F, G**).

We further evaluated cognition with NOR and OCD with marble burying test. For NOR we applied discriminatory index (DI) calculated as (total time investigating novel object - total time investigating familiar object) / total time investigating both objects. The DI showed significant difference between WT and *mdx52* mice suggesting impaired memory retention in *mdx52* (*P* = 0.0312). Significant improvement in DI was observed only following IV delivery (*P* < 0.0001) and no effect was observed with ICV (**Fig. 2H**).

We explored marble burying test, representing anxiety and OCD in *mdx52*; this test was normalised to grip strength, as this task involves motor co-ordination with grip and the muscle disease is not treated with neuronal targeted gene therapy. We observed significant decrease in % unhidden marbles in *mdx52* suggesting anxiety-OCD behaviour (*P* = 0.0086) with no effect of ICV treatment; surprisingly we observed normalisation to WT (*P* = 0.0256) following IV delivery (**Fig. 2I**). There was no significant difference with marble burying when compared directly, without grip strength normalisation (**Supplementary Fig.1B**).

Overall, these results demonstrated improvement with both delivery methods in emotional reactivity assessments. A surprising observation is the superior efficacy with IV delivery in elevated zero maze and efficacy only with IV delivery for NOR and marble burying.

### IV delivery results in widespread brain transduction with improved targeting of hindbrain and cerebellum compared to ICV

To better understand the superior efficacy observed with IV delivery, we undertook detailed μDys /dystrophin expression analysis in the brain with immunofluorescence, Simple western (WES), transcripts and VGC (**Fig. 3, Supplementary Fig. 2C,D**).

**Figure 3.**
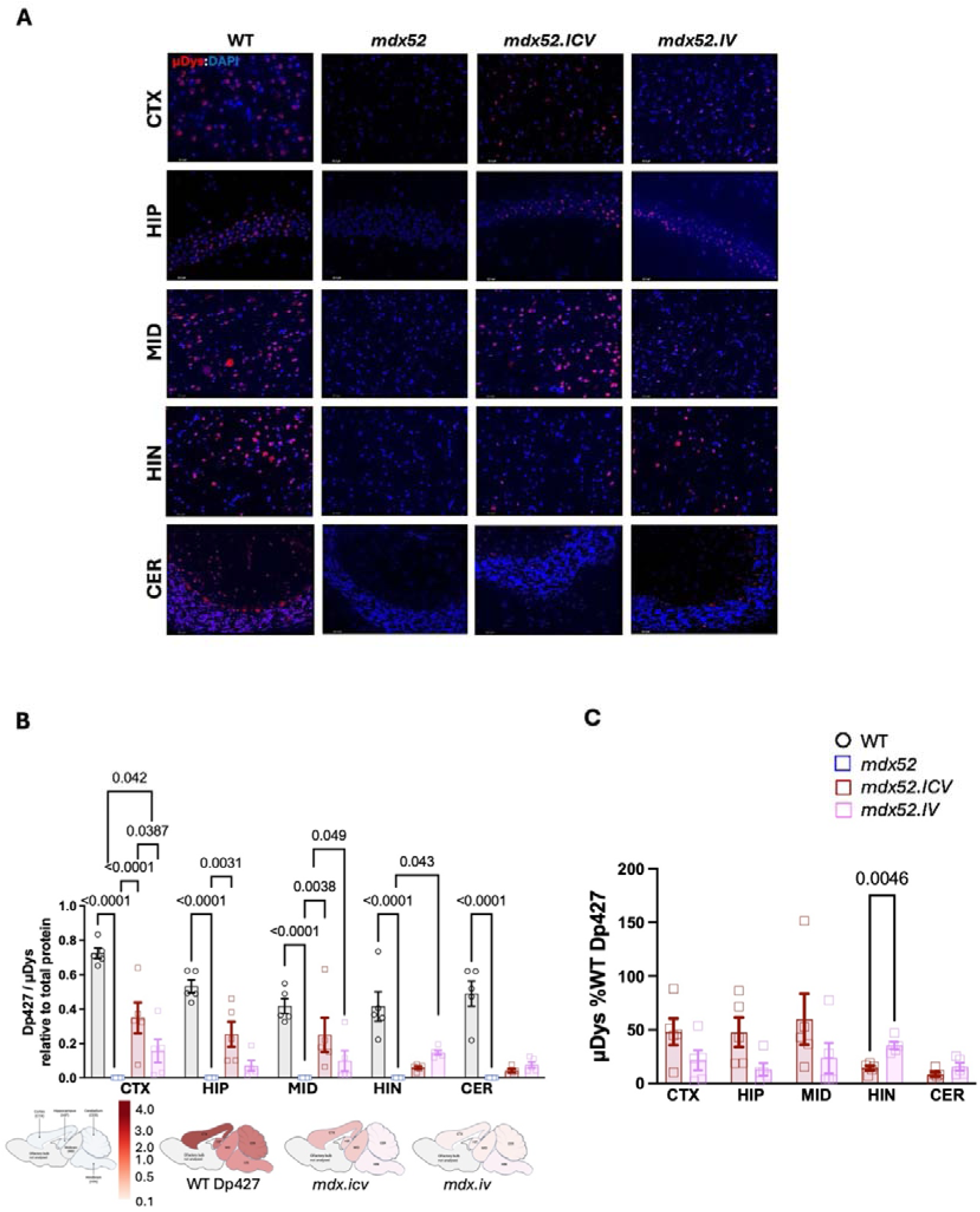
IV delivery results in widespread brain transduction with improved targeting of hindbrain and cerebellum compared to ICV. Brains collected 12 weeks post-gene therapy and dissected into cortex (CTX), hippocampus (HIP), midbrain (MID), hindbrain (HIN), and cerebellum (CER) **(A)** Immunofluorescent staining for dystrophin/μDys (NCL-DYS3, Leica Biosystems) in WT, *mdx52*, *mdx52.ICV* and *mdx52.IV* **(B)** Dp427 or μDys protein relative to total protein by simple western capillary (WES) (n=5 per group). **(C)** μDys as %WT Dp427 (n=5 per group). Data presented as mean ±SEM. Two-way ANOVA with Tukey’s multiple comparison test (p-value<0.05).

Sagittal whole brain immunofluorescence for endogenous dystrophin in WT or μDys (NYS-DYS3/ Hinge 1) showed positive staining in cortex, hippocampus, midbrain and cerebellum in WT and both groups of treated *mdx52*. There was no dystrophin staining in control *mdx52*. With ICV delivery there was higher μDys staining in the fore to midbrain regions and less in hindbrain and cerebellum, in-keeping with AAV9 ICV rostrocaudal biodistribution. With IV delivery there was generally lower levels of μDys staining but more uniform expression throughout the brain (**Fig. 3A**). Dp427 or μDys on WES affirmed the brain biodistribution profiles observed by immunofluorescence, with slightly higher expression in hindbrain and cerebellum with IV delivery (**Fig. 3B**). Neither delivery method achieved WT endogenous levels of Dp427. For purposes of comparison to brain delivered ASO strategies we also quantified as %WT Dp427 (**Fig. 3C**, **Supplementary Fig. 2 %WT Dp140**). Overall, these results suggest IV delivery achieved lower expression levels but wider-spread transduction with improved targeting of hindbrain and cerebellum compared to ICV.

### Neuronal targeted **μ**Dys restores post-synaptic DAPC interactors and known functional interacting complexes associated with Dp427 and Dp140 isoforms

DAPC have important roles in the CNS and in neurons, where they participate in the postsynaptic clustering and stabilisation of inhibitory GABAergic synapses. We evaluated the effects of AAV.μDys on DAPC and indirect functional complexes, neuronal post-synaptic gephyrin, GABA_A_R^55,56^ and pre-synaptic Dp140 associated vesicular glutamate transporter (VGLUT1).^26^ (**Fig. 4A, 5A**).

**Figure 4.**
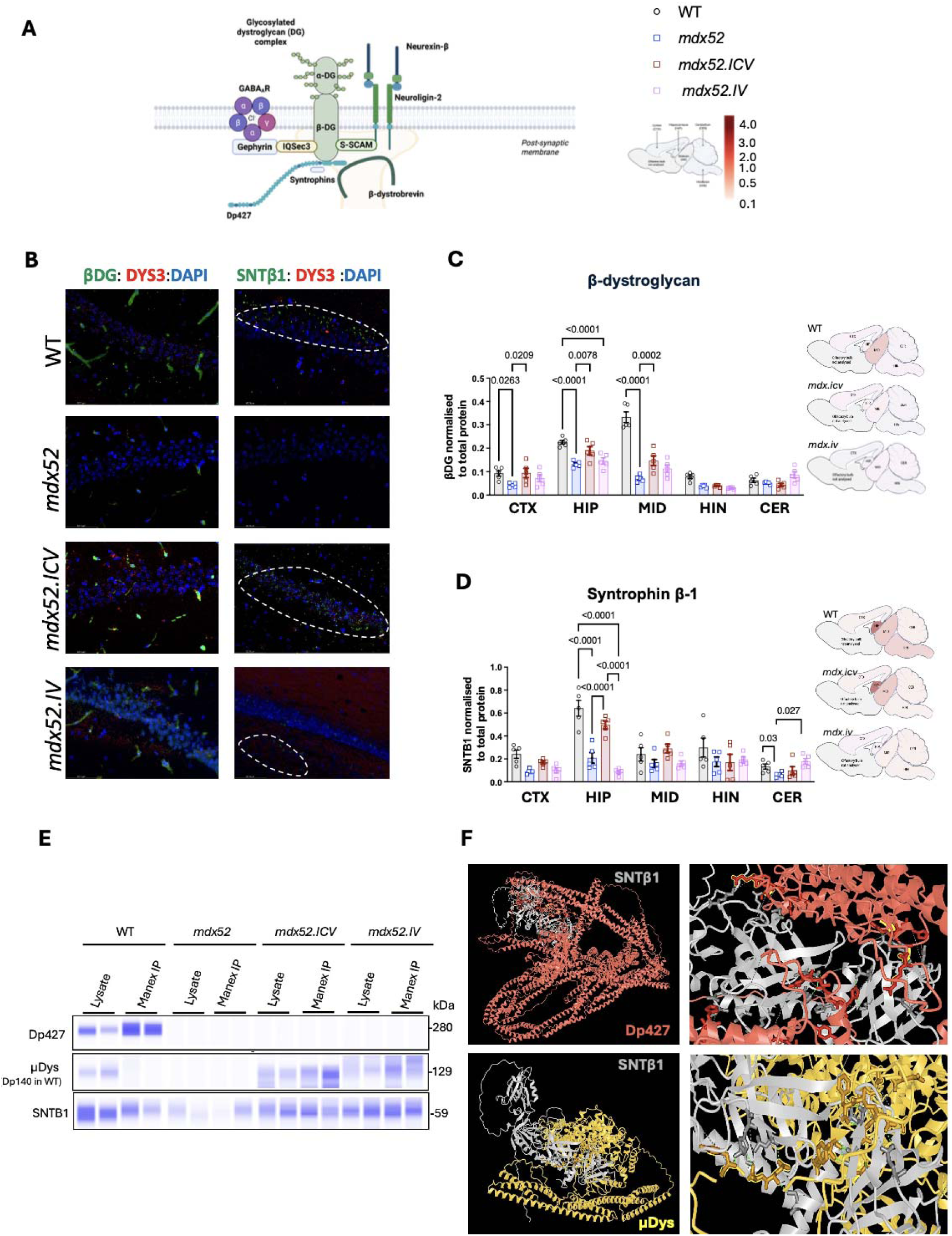
Neuronal targeted μDys restores post-synaptic DAPC interactors. **(A)** Schematic of post-synaptic neuronal DAPC and brain region expression schematic. (**B**) Double labelled immunofluorescent staining of hippocampal regions for Dp427/μDys with Beta dystroglycan (ßDG) or syntrophin beta 1 (SNT1ß). (**C**) ßDG brain regional expression by simple western normalized to total protein (**D**) SNT1ß brain regional expression by simple western normalized to total protein (**E**) Co-immunoprecipitation to Dp427/ μDys with Syntrophin beta 1 (**F)** Protein-Protein interaction modelling of SNTß1 (grey)with Dp427 (red) and Dys (yellow), interaction at ABD and Hinge 1 enlarged on right panel. Figures are created with AlphaFoldserver.com Groups size n= 5 per group and data is mean ± SEM. Two-way ANOVA with Tukey’s multiple comparison test (p-value<0.05).

**Figure 5.**
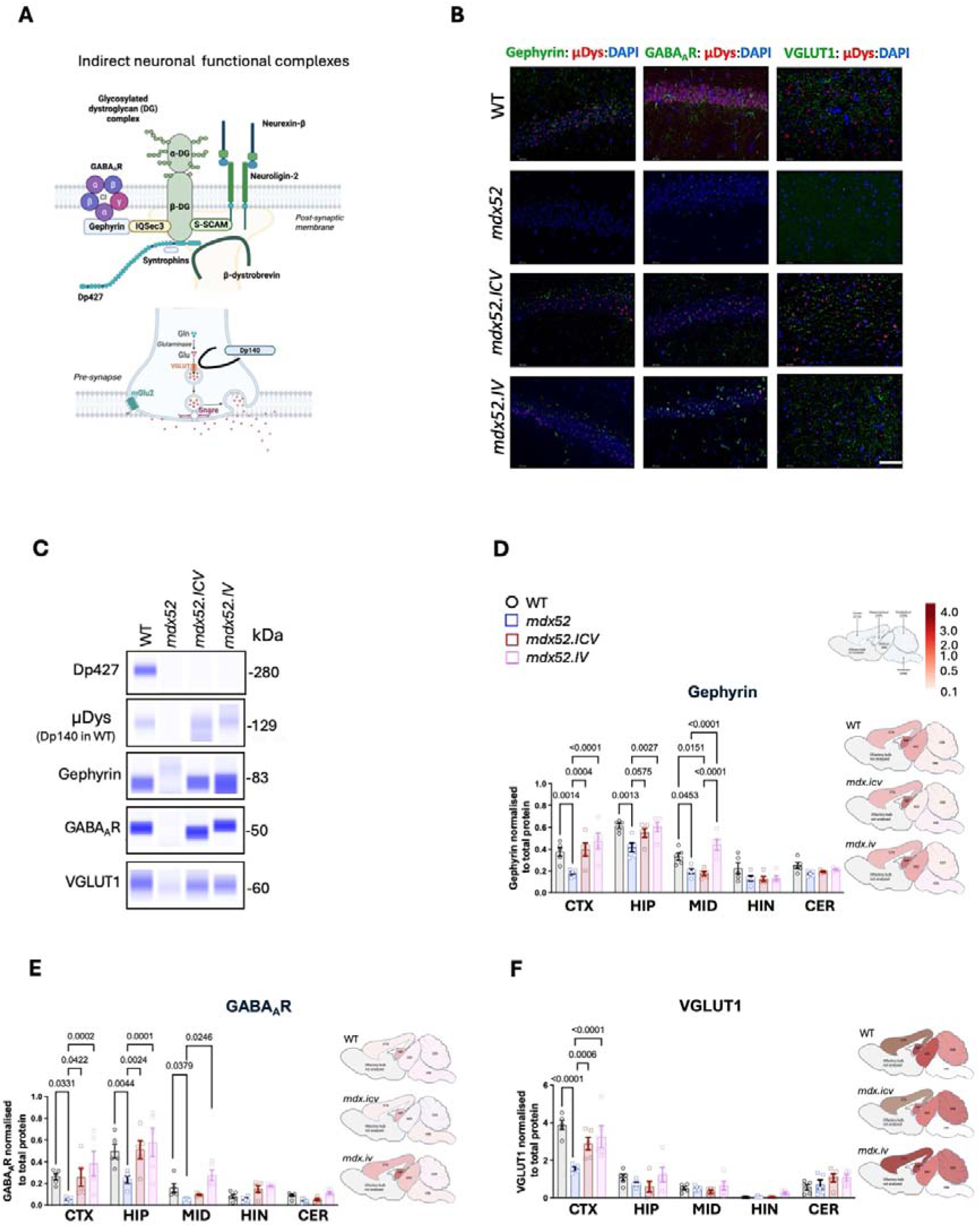
Neuronal targeted μDys restores synaptic indirect functional complexes. **(A)** Schematic of post-synaptic indirect Dp427 function complexes gephyrin and GABA_A_R and presynaptic indirect Dp140 functional complex VGLUT1. Figure created with Biorender.com (**B**) Representative co-immunoprecipitation for dystrophin or μDys with associated proteins demonstrated on WES n=5. (**C**) Double labelled immunofluorescence in hippocampus for dystrophin or μDys and functional interactor complex. Brain regional expression by simple western normalized to total protein for **(D)** Gephyrin (**E**) GABA_A_R (F) VGLUT1. (n=5 per group) mean ± SEM and analysed Two-way ANOVA and Tukey’s multiple comparison test-hoc.

For DAPC restoration immunofluorescence for dystrophin/μDys with Beta-dystroglycan (βGD) and Syntrophin beta 1 (SNTβ1) showed positive staining in hippocampus in WT and absence or reduction in *mdx52*. Both ICV and IV delivery restored βGD hippocampal staining, whilst SNTβ1 positive staining was observed with both delivery methods, the hippocampal biodistribution was only restored with ICV delivery (**Fig. 4B**).

On WES, βGD was significantly reduced in *mdx52* cortex, hippocampus and midbrain (*P =* 0.0263*, P* < 0.0001*, P* < 0.0001 respectively), with no significant cerebellar difference. Following ICV delivery, βDG levels were restored to WT levels; cortex (*P* = 0.0263, 98%WT), hippocampus (*P* < 0.0001, 83%WT) and midbrain (*P* < 0.0001, 44%WT). With IV delivery there were non-significant increases in cortex and midbrain, reflecting the immunofluorescence and lower μDys expression with IV (**Fig. 4B, C**).

SNTβ1 was significantly reduced in the hippocampus and cerebellum in *mdx52* (*P* < 0.0001*, P* = 0.0300 respectively) and restored to WT levels following ICV delivery in hippocampus (*P <* 0.0001, 77%WT) whilst IV restored cerebellar levels only (*P* = 0.0270, 99%WT) (**Fig. 4D**). Syntrophins interaction with Dp427 are characterised with muscle isoform to be rods 17,22 and CTD,^57,58^ but the protein-protein interactions in brain dystrophin and μDys is not well documented. We affirmed interaction of SNT1β to Dp427 in WT and in μDys treated *mdx52* with co-immunoprecipitation to Dp427/μDys Hinge 1 (**Fig. 4E**). We further modelled interacting domains *in silico* with alphafold 3 and demonstrated known protein-protein interactions at C terminal domain (CTD) between Dp427 and SNTβ and modelled a novel putative interaction with PDZ domain of SNTβ1 and common region at actin binding domain (ABD) and Hinge 1 in Dp427 and μDys. (**Fig. 4F, Supplementary Fig. 3**)

Following on we assessed restoration of synaptic dystrophin associated functional complexes gephyrin, GABA_A_R and VGLUT1 and demonstrated association of these proteins with Dp427 and μDys with co-immunoprecipitation, that were absent in control *mdx52* (**Fig. 5A, B)** On immunofluorescence we observed absence of those proteins in *mdx52* mice and restoration in ICV and IV treated animals (**Fig. 5C**).

Gephyrin is a postsynaptic scaffold protein, linked to GABA_A_R clustering, mediated by indirect interactions with brain Dp427; Gephyrin was significantly reduced in cortex, hippocampus and midbrain of *mdx52* (*P* = 0.001, *P* = 0.0013*, P* = 0.0453 respectively). Following ICV delivery there was restoration to WT levels in cortex only (*P* = 0.0004, 99%WT) with trend in hippocampus, with no effect on midbrain. In contrast IV delivery was superior with restoration of gephyrin equivalent to WT profile (cortex *P* < 0.0001, 99%WT, hippocampus *P* = 0.0027 98%WT, midbrain *P* < 0.0001, 99%WT) (**Fig. 5C, D**).

Associated with gephyrin reduction is the mislocalisation of post-synaptic GABA_A_R in *mdx*.^24,55,59–61^ We confirmed significant reduction in cortex, hippocampus and midbrain (*P =* 0.0331*, P* = 0.0044*, P* = 0.0379). With ICV, levels were only restored in cortex and hippocampus (*P* = 0.0422, 97%WT, *P* = 0.0024, 102%WT respectively) and not midbrain. Whilst again we observed restoration to WT profile with IV delivery (cortex *P* = 0.0002,101%WT, hippocampus *P* = 0.0001,103%WT, and midbrain *P* = 0.0246, 99%WT) (**Fig. 5E**).

Dp140 deficiency is associated with altered glutamatergic neurotransmission, reduced presynaptic VGLUT1 levels, and impaired docking of synaptic vesicles in basal lateral amydala.^62^ We observed significantly reduced cortical VGLUT1 in *mdx52* mice (*P < 0.0001*) where VGLUT1 is highly expressed.^63^ Following gene therapy we observed significant cortical increase of VGLUT1 (ICV *P* = 0.0006 74%WT, IV *P* < 0.0001, 84%WT) while no genotypic differences were observed in the other brain regions analysed (**Fig. 5F**).

Overall, we observed restoration to WT levels of βDG with ICV delivery and regional restoration for SNTß1 dependent on delivery modality. Again, we observe superior restoration of synaptic functional complexes gephyrin and GABA_A_R with IV delivery and equal effect on cortical VGLUT1 irrespective of delivery routes.

### Intravenous neuronal targeted AAV9.**μ**Dys restores neuromuscular junction function, grip strength and motor coordination

The most striking finding was the significant improvement in grip strength exclusively noticed with neuronal targeted AAV9.μDys delivered intravenously to *mdx52* (*P* = 0.03) whilst there was no effect in ICV delivery (**Fig. 6A**). We noted very low VGC in muscle and heart of IV-treated *mdx52* but no μDys protein was detected in either treated groups, due to transcriptional neuronal selectivity of hSyn promoter (**Fig. 6B-D**).

**Figure 6.**
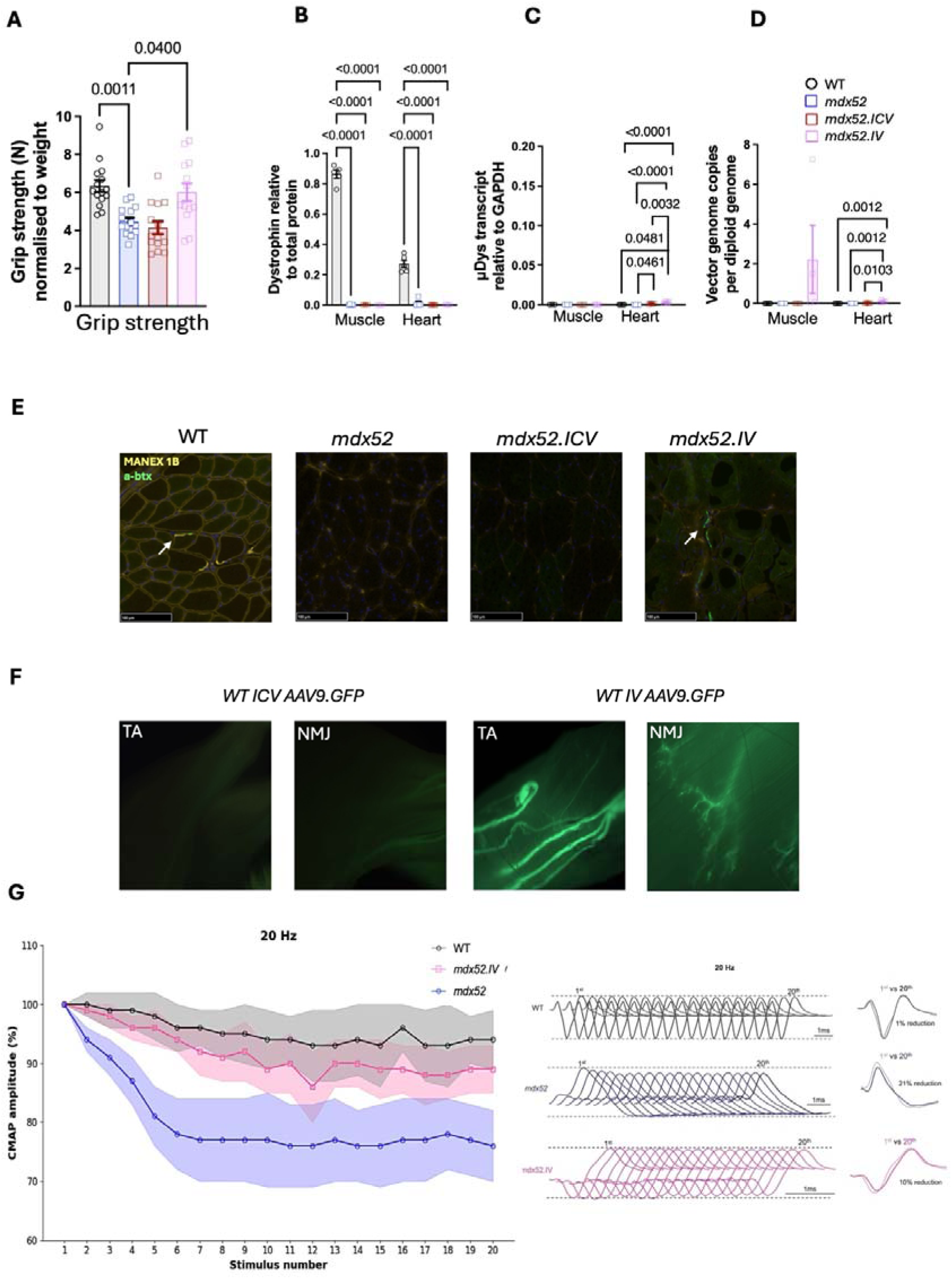
IV delivered AAV neuronal targeted μDys expressed in neuromuscular junction restoring grips strength and electrophysiological deficit. **(A)** Grip strength (n=14-15 per group) **(B)** Dystrophin/ μDys expression in Tibialis Anterior (TA) muscle and heart by **(B)** WES, **(C)** μDys transcripts and **(D)** μDys VGC (n=4-5 per group, TaqMan Multiplex qPCR to mouse GAPDH) **(E)** Immunofluorescence of TA depicting neuromuscular junction labelled with α-Bungarotoxin (green) and dystrophin/ μDys with MANEX1B (yellow). Inserts: 100um **(F)** NMJ transduction after AAV9.hSyn.GFP vector delivered to P1 WT pups ICV or IV (n=4 per group). Stereoscopic fluorescence images taken of muscle at 4 weeks post-delivery **(G)** CMAP amplitude as percentage of 1^st^ pulse after RNS at 20Hz (left). Example of RNS at 20Hz (right) representing the reduction in WT (black), *mdx52* (blue), *mdx52.IV* (pink). Data are presented as mean ± SEM, one-way ANOVA, followed by Dunnet’s test (p-value<0.05).

To identify further mechanistic insights to this effect, we thoroughly assessed muscle immunohistochemistry for off-target transgene expression at the sarcolemma in TA.

When analysing the TA muscle sections of the IV μDys treated *mdx52* mice we affirmed no dystrophin positive myofibres in treated *mdx52*. However, we identified clear dystrophin immunohistochemical staining at structures morphologically suggestive of neuromuscular junctions (NMJ). We firstly confirmed that in WT mice the Dp427m isoform (recognisable by the antibody MANEX1B directed against the unique first exon) colocalised with bungarotoxin in WT, and as expected saw no co-staining in control *mdx52* or ICV treated *mdx52* (**Fig. 6E**). Subsequently we demonstrated clear colocalization between the μDys and bungarotoxin, suggesting that the transgene was expressed at the NMJ in *mdx52* treated IV. To further illustrate peripheral nervous system transduction differences between ICV and IV delivery we generated an AAV9.hSyn.GFP equivalent vector delivered to P1 WT and stereoscopic fluorescence images of muscle at 8 weeks demonstrate clearly only IV delivery transduced the NMJ and differences between brain transduction (**Fig. 6F, Supplementary Fig 4**).

To test the effect of AAV9.hSyn.μDys in restoring NMJ function, we performed repetitive nerve stimulation (RNS) to study neuromuscular transmission, which previously demonstrated post-synaptic NMJ defects in *mdx* mice.^64^ We implanted the TA with fine-wire intramuscular electrodes, and through peripheral nerve stimulation (sciatic nerve) supramaximal stimuli were delivered at 3, 10, 20 and 40 Hz (20 pulses per train) and the change in compound muscle action potential (CMAP) amplitude was quantified as the ratio of the last to the first response (**Supplementary Fig.5A**). WT animals showed minimal CMAP decrement across frequencies, consistent with intact NMJ transmission (**Fig. 6G**). In contrast, *mdx52* mice exhibited pronounced decremental responses at both low and high frequencies, with progressive reductions in CMAP amplitude across the stimulus train. Notably, *mdx52.IV* animals displayed only minimal CMAP decrements comparable to WT across all frequencies, indicating that restoration of dystrophin expression at the NMJ functionally rescued neuromuscular transmission. Representative response and traces for 20 Hz are shown in **Fig. 6G**. Therefore, we demonstrated that the μDys was expressed at the NMJ in IV treated *mdx52* leading to functional restoration of CMAPs.

To further explore the *in vivo* functional effects of NMJ μDys expression we undertook further motor co-ordination tests. The beam walk is used to assess motor coordination with significant difference between control WT and *mdx52* (Beam Scor*e, P* = 0.0001, Time*, P =* 0.0247) and again superior IV efficacy was observed restoring beam walk to WT equivalent (Score*, P* = 0.043, Time *,P =* 0.0361), whilst there was no significant effect on either measure with ICV (**Fig. 7A, B**).

**Figure 7.**
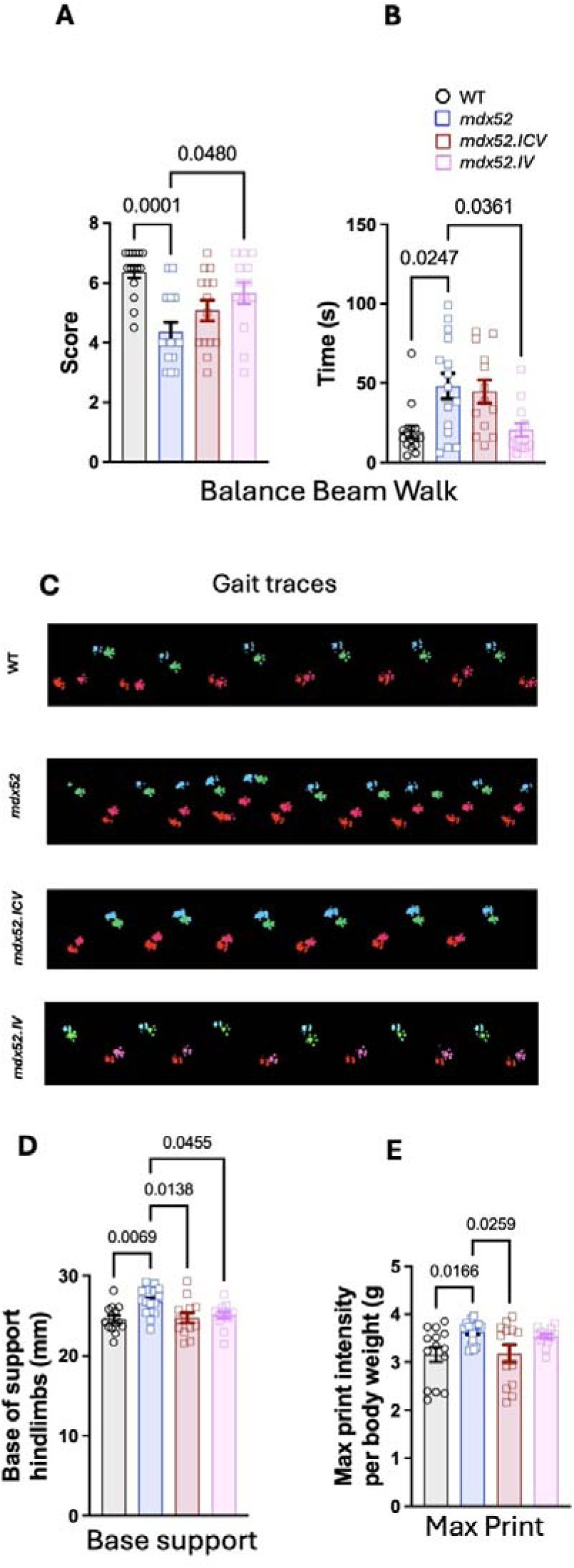
IV AAV9.μDys gene therapy improves motor coordination. Beam walk balance test (**A**) score (**B**) Time to traverse 6mm beam **(C)** Representative Gait traces of paw prints from CatWalk XT (Green: Left-Front, Pink: Right-Front, Red: Right-Hind and Light blue: Left-Hind paws) at 12 weeks old **(D)** Base of support in hindlimbs **(E)** Max print intensity (n= 14-15 per group). Data presented as mean ± SEM and analysed with one-way ANOVA with Tukey’s post-hoc.

CatWalk XT was used for analysing gait and illustrated significant differences between WT and *mdx52* in several parameters (**Supplementary Fig. 5, Supplementary Table 2**). Following treatment improvement was observed in some gait parameters in treated *mdx52* as illustrated by gait traces (**Fig. 7C**, **Supplementary videos 1-4**). In particular gait stability and coordination improved with both modalities demonstrated by base-support, and weight bearing improved with ICV only as maximum print intensity (**Fig. 7 D, E**). There was no effect on rotarod latency to fall with either treatment group suggesting treatment for muscle disease maybe the more significant component for this test (Supplementary 5E).

In summary we demonstrated that IV delivery resulted in NMJ transgene expression (without myofibre expression) with restoration of post-synaptic CMAPs defect, and *in vivo* functional effects with normalisation of grip strength and beam walk. There was mixed effect with more complex motor assessments that likely involve both central, peripheral and muscle components with gait and rotarod studies (**Supplementary Fig. 5**).

## Discussion

The neurocognitive and neurobehavioural co-morbidities in DMD are under-recognised, representing a significant unmet clinical need, not addressed by current therapies. Recent studies in the *mdx* and *mdx52* mouse confirmed similar brain involvement and the potential for its progression over time.^26,53^ As dystrophin is lacking in muscle, heart and brain in patients with DMD, it would be ideal to treat all organs lacking dystrophin, so to avoid unmasking, in the longer term, the potential for progressive CNS dysfunction, as observed in peripherally treated Pompe disease patients.^65^

Our study is the first to evaluate the role of neuronal targeted AAV.μDys mediated gene supplementation as therapeutic approach for brain dystrophin deficiency in an animal model of DMD, and to demonstrate significant neurobehavioural improvement, often to WT levels, depending on the behavioural tasks assessed, the mode of AAV administration and resulting AAV biodistribution. Unexpectedly, the neuronal promoter driven µDys was also associated with improvement in motor functional abilities and grip strength with important implication for the development of effective therapies for DMD.

µDys optimised for protection of muscle from contraction induced damage and under the control of muscle specific promoters are in clinical trials with one of them, ELEVIDYS, received FDA approval for ambulant DMD patients in 2023. While µDys cannot recapitulate the complex pattern of expression of multiple dystrophin isoforms in the brain, the question we intended to address was if the neuronally expressed µDys containing exons 1-17 and 59-71, including hinges 1, 2 and 4 (**Fig. 1B),** was able to provide functional benefit and restore functional complexes relevant for brain function.

One essential question for brain-related translational research is whether postnatal restoration could provide some beneficial effect. This had been previously addressed using direct delivery of ASOs to the brain via osmotic pumps, intrathecal or via the cisterna magna.^37,40,66–68^ Multiple studies using ASOs have demonstrated partial improvement of behavioural aspects and restoration of fear response^38–40,69,70^ although the levels of dystrophin restoration in the brain were low across those studies.

Comparative analysis of brain-delivered splice switching ASOs with different backbone chemistries and administration routes revealed limited biodistribution, partial efficacy and dose-limiting toxicity, highlighting one of the challenges for CNS-targeted ASO therapies.^40^ These limitations motivated us to explore AAV gene therapy to deliver a micro-dystrophin to the brain of *mdx52* mice. In view of the well demonstrated role of dystrophin in neuronal GABAergic and glutamatergic transmission, we used a synapsin promoter to drive μDys expression exclusively in neurons. Both the postnatal IV and ICV delivery of AAV9.hSyn.µDys robustly demonstrated amelioration of emotional reactivity, abnormal fear and anxiety in multiple behavioural assessments in *mdx52* irrespective of delivery route (**Fig. 2**). In some of these tests we observed complete normalisation of the characteristic behavioural features observed in the *mdx52*, with expression in cortex, hippocampus and midbrain as common regions transduced by both delivery methods with AAV9 (**Fig. 3**).

While both IV and ICV were effective in most of the behavioural tasks studied, some differences were noted. We observed higher efficacy with IV delivery on elevated zero maze distance in open arms, learning and OCD behaviours (**Fig. 2**). NOR test has shown differences in DMD models but there is a variable phenotype across *mdx23, mdx5cv, mdx52*.^52,71^ Previously ICV delivery of tricyclo-DNA ASO demonstrated improvement on object recognition task.^66^

Our gene therapy demonstrated normalisation of the discrimination index (DI) in the object recognition test between *mdx52* and WT following IV delivery but not ICV. The brain regions involved in recognition memory are perirhinal cortex, hippocampal CA1 region and medial prefrontal cortex^72^ and hindbrain, linked to object recognition.^73^ Although expression of µDys was higher in cortex and hippocampus in ICV, the lower levels of expression in hindbrain compared to IV may have resulted in lack of effect, demonstrating the advantage of a broader biodistribution profile to determine efficacy outcomes for NOR (**Fig. 2H, 3C**) .

For the first time in a *mdx* mouse model, we assessed marble burying, as a measure of OCD behaviour. When normalised to grip strength, marble burying was restored to WT levels with IV only, while no effect with ICV. OCD, a well-recognised comorbidity of DMD boys, is associated with imbalance in glutamate and GABAergic activity in cortico-striato-thalamo-cortical circuits^74^; following neonatal AAV delivery using both administration routes, these brain regions showed robust μDys expression also confirmed clearly by AAV9.GFP study (**Supplementary Fig. 4**). The superior efficacy with IV may reflect a more uniform lower expression levels across the neural circuits and transduction of hindbrain and cerebellum. The rostrocaudal transduction gradient achieved with ICV delivery results in higher forebrain expression, but fails to transduce the hindbrain and cerebellar regions.

The improved efficacy in restoration of multiple emotional reactivity behaviours after AAV gene therapy compared to ASO likely relates to the differing biodistribution and higher levels of μDys expressed (range 9-69%WT Dp427), achieving higher levels compared than previously achieved with ASOs, providing evidence of clear superiority of the AAV gene supplementation in the *mdx* mouse model (**Fig. 3C**). Our data also clearly outperforms the recently reported findings on ICV and cisterna magna administration of AAV9-U7 providing constitutive ASO expression to skip exon 51 in the *mdx52*. Surprisingly this study failed to improve the emotional behaviour of adult *mdx52* despite achieving levels of Dp427 restoration between 6-12%.^68^ This is surprising as previous studies delivering ASO to restore Dp427 in adult *mdx* mice resulted in behavioural improvement.^38,39^ The inability of exon 51 skipping to restore Dp140 expression might be a contributory factor as the translational initiation codon of this isoform is in exon 51^37,38^ The 3’ end mRNA depletion present in both the muscle and brain of *mdx* mice, linked to both transcript imbalance, and epigenetic changes at the DMD locus, might have also contributed to the lack of clinical efficacy.^75^

To further underpin mechanistic functional impact of neuronal μDys expression we assessed for the first time in a single study the effect of the AAV gene therapy on several components of DAPC and indirect functional complexes which are depleted in *mdx52* brain namely ßDG, SNT1ß, gephyrin, GABA_A_R and VGLUT1.

βDG interacts with Dp427c and Dp427p at the neuronal post-synaptic densities of excitatory and inhibitory synapses and glial expressed Dp140 at the CRD.^76^ It is highly expressed in hippocampal pyramidal neurons and cerebellar Purkinje neurons, where it plays a versatile role in cell signalling related to cell polarity.^77,78^ We observed significant reduction in cortex, hippocampus and midbrain with ßDG in *mdx52* that was restored with ICV and IV AAV9.µDys as it expresses the CRD interacting domain in neurons.

Other components of DAPC include syntrophins and we showed for the first time significant reduction in SNTβ1 in hippocampus and cerebellum between control WT and *mdx52* with restoration to WT levels with ICV delivery in hippocampus and restoration in cerebellum with IV delivery. The syntrophin interacting domains with Dp427 in muscle are well delineated at Rods 17, 22 and CTD^57,58^ that are not included in this µDys transgene. There is limited characterisation of SNTβ1 with brain dystrophins and through co-immunoprecipitation we demonstrate interaction with Dp427 and µDys with putative common novel interacting domain in ABD and Hinge 1 predicted with protein-protein modelling (**Fig. 4**).^57,58^

In assessment of effect on indirect functional interacting complexes we observed superior efficacy with IV delivery at restoring the brain profile of gephyrin and GABA_A_Rs (**Fig. 5**). (**Fig. 2**). Gephyrin is a microtubule-associated protein interacting with Dp427^60^, involved in membrane protein-cytoskeleton interactions and clusters with GABA_A_Rs.^79,80^ GABA_A_Rs do not interact directly with dystrophin, with likely intermediate interactions through synArfGEF, which binds to gephyrin.^23,81^ The mislocalisation of GABA_A_ receptor subunit, α2, in absence of Dp427 at the inhibitory synapse contribute GABAergic dysfunction resulting in fear and anxiety features which characterise the mouse model and patients with DMD.^24,56,59,60,82^ The μDys levels were higher in ICV than IV delivery but interactor restoration, suggests post-natal equivalent of WT levels of dystrophin are not required to restore emotional reactivity behaviours and are more related to having sufficient level of interactor restoration.

In this comprehensive study we also assessed for VGLUT1 protein restoration that has direct effect on glutamatergic neurotransmission. Previous studies show absence of VGLUT1 in basal lateral amygdala (BLA) *in mdx52* and restoration of VGLUT1 levels and localisation with direction infusion with ASO to BLA in relation to social behaviour.^26^ We did not observe a hippocampal difference but did not specifically dissect the BLA for study; nevertheless we restored significantly reduced cortical VGLUT1 with ICV and IV delivery in *mdx52* mice. VGLUT1 is responsible for packaging the excitatory neurotransmitter glutamate into synaptic vesicles for release and is most highly expressed in cortex.^83^ It plays a role in synaptic plasticity, memory and cognitive processing and is reduced in the cortical neurons in dementia.^84^ The implications of VGLUT1 deficiency in DMD are likely widespread with potential effects on cognitive, social and memory deficits. Our restoration of cortical VGLUT1 in combination with normalisation of gephyrin and GABA_A_R in all affected brain regions (cortex, hippocampus, midbrain) was only achieved with IV delivery. The biodistribution achieved with IV may explain the superior efficacy compared to ICV and also highlights that µDys is able to impact excitatory VGLUT1 and inhibitory GABAergic dysfunction .

A very striking finding of our study is the normalisation of grip strength with neuronal targeted µDys and expression in the NMJ (**Fig. 6**). Several studies have reported the NMJ abnormalities in DMD mouse models, which compound muscle weakness and progressive NMJ degeneration^85–88^. These structural and functional NMJ defects likely contribute to the impaired motor output observed in *mdx* mice and may therefore influence the grip strength improvements observed following gene therapy. In keeping with the NMJ localisation of μDys and improved strength, we also demonstrated improvement of the *mdx*52 neuromuscular transmission defect following gene therapy (**Fig. 6F**). Similar although only partial results were obtained when delivering antisense oligonucleotide therapy on *mdx23* mouse model were limited improvement of NMJ function was observed.^64^

While the relationship between the deficiency of different brain isoforms and brain function is well recognised, recent data have also reported that brain dystrophin deficiency is also associated with lower peak of functional activity and grip strength^89^ in DMD individuals.^30,90^ These findings are replicated in *mdx52* with reduced grip strength compared to *mdx23*, suggesting an elusive role of brain dystrophin isoforms in motor function.^30^ This adds to the central component of cerebellar dystrophin deficiency which has also been associated with reduced motor coordination^91^. The result of our multimodal assessment of motor function and strength clearly indicated that neuronal driven µDys was associated with improved coordination, muscle power, as demonstrated by normalisation balance beam walk with only IV delivery (**Fig. 7A, B**). The CatWalk XT gait analysis showed improvement in gait stability and coordination with both delivery routes, weight bearing with ICV delivery only (**Fig. 7C-E, Supplementary gait videos 1-4**) but there was no effect on several other motor parameters (**Supplementary Fig. 5**, **Supplementary Table 2**) that may relate to neuronal targeted gene therapy.

In summary our study is the first to apply AAV µDys gene supplementation to address comorbidities associated with Dp427 and Dp140 brain dystrophin deficiency in *mdx52* mice with 2 delivery modes. Our approach ameliorated multiple neurobehavioural emotional reactivity behaviours with neuronal targeted expression with demonstration for the first time effects on both pre- and post-synaptic brain dystrophin interactors and associated complexes involved in GABA and glutamatergic neurotransmission in a single study. These results are highly relevant to the behavioural comorbidities that characterise patients with DMD.

The breadth of gene therapy efficacy was contingent on delivery method and biodistribution profile achieved with associated expression in NMJ with IV delivery associated with further improvements in cognition, OCD, motor coordination and grip strength. It is also important to note that in our studies we delivered μDys to newborn mice, whilst previous splice switching ASO studies in adult *mdx* mice had clearly demonstrated partial behavioural improvement, suggesting that further studies treating adult mice will be needed before considering our approach for its potential clinical translation.

Nevertheless, this is the first study to demonstrate a clear and robust improvement of a wide range of behavioural abnormalities in the *mdx52* mice and restoration of the DAPC complexes which bodes well for future studies in this area. Our findings on the improved NMJ junction transmission and function after neuronal targeted μDys expression adds value to our novel approach which could contribute to clinical translational application.

## Data Availability

The data that support the findings of this study are available from the corresponding author, upon reasonable request.

## Acknowledgements

We are grateful to Ayana Withana for performing protein extractions, Jenny Vartiainen qPCRs, Camila Vallve behavioural assessments and Riccardo Privolizzi ICV injections. We also thank Dr Jun Tanihata and Dr Shin’ichi Takeda (both National Center of Neurology and Psychiatry, Tokyo, Japan) for providing the *mdx52* mouse breeders. We are thankful to BIND consortium for establishing the battery of behavioural assessments used on this study and Muscular Dystrophy UK for covering the salary of researchers. We are grateful to the UCL Animal Research Facility, for mouse breeding, care, and genotyping.

