## Supplementary material for "Neuronal targeted AAV Micro-Dystrophin restores neurobehavioural co-morbidities, grip strength and motor coordination in *mdx52* mouse model of Duchenne Muscular Dystrophy"

##### **Supplementary Materials and Methods**

##### **Emotional Reactivity**

##### Freezing Response

Mice were restrained by the same experimenter by grasping the scruff and back skin between thumb and index fingers, while putting the tail between the third and little fingers. The animals were then tilted upside-down in a position that the ventral part of their body faced the experimenter. After 15 seconds, the mice were released into a novel cage containing clean sawdust and then video-tracked for 5 min using SMARTv3.0 software (Panlab, Harvard Apparatus). The tonic immobility was quantified during this period of time, since this is the characteristic of unconditioned fear responses induced by this short acute stress. Freezing response was identified as complete immobilization of the mouse, except for respiration.^89^ The amount of tonic immobility was expressed as the percent time spent freezing and calculated for group comparisons.

##### Light Dark Choice

The light/dark choice test utilises a box with two equally sized compartments at 20x40x35cm (width x length x height) (Stoeling Europe). One compartment is covered and black, and another open and brightly illuminated, connected via a small opening door. To visualise the animal’s movements, the dark compartment is illuminated with an infrared illuminator at 850nm (Univivi) and captured using an infrared camera (Computer TG4Z2813FCS-IR 1/3" 2.8-12mm) positioned overhead. Each mouse was placed in the dark compartment for 10 seconds before the door was opened, and the mouse was allowed to explore freely the whole apparatus for 5 minutes. Number of entries and total time spent in the lit compartment were analysed using ANY-maze software 7.40 (Stoelting Europe).

##### Open Field

Open field exploratory behaviour (25cmx25cmx25cm) was recorded for 30 minutes and data analysed using SMARTv3.0 software (Panlab, Harvard Apparatus). The total distance travelled, distance spent in the center and the mean speed in the total area were calculated.

##### Elevated Zero Maze

The apparatus was an elevated ring (zero) shaped maze (60cm in diameter, 5cm wide track, 16cm high walls, elevated 62cm above the floor, Ugo Basile, Gemonio, Italy), composed of two open (unwalled) and two closed (walled) quadrants of equal size. Overhead lighting provided 500-600 lux in open arms, while closed ones were luminated at 30-40 lux. The test was performed for 5 minutes using a ceiling-mounted camera and automated tracking was performed with SMARTv3.0 software (Panlab, Harvard Apparatus). The mice were firstly placed in the centre of the open quadrant and allowed to explore freely. The mice nature makes them lean toward darkly lit area to stay hidden from the sight of predators; the lesser time they spend in the darkly lit segments are deemed to have lower levels of anxiety. The number of entries and time spent in open or closed arms were counted for group comparisons.

**Cognitive and Recall**

### Novel Object Recognition Test

Two days before testing, the mice were habituated with the test environment by placing them in an empty open-field chamber for ten minutes. One day before the test, the mice were familiarised with two identical small objects by placing them in the chamber with the objects for ten minutes. The mice were tested by placing them in the chamber with one of the original objects and one novel object for five minutes. Their movements were filmed by a camera and analysed by ANY-maze software to quantify the time each mouse spent exploring each object.

**OCD behaviour**

##### Marble Test

The marble test is indicative of repetitive and compulsive behaviours which relates to OCD.^90^ Mice are known to make burrows and dig in natural environment for their own protection against any harm. The test was assessed by placing twenty marbles in five rows and counting the number of marbles buried after five minutes. This test has not been used to assess DMD brain deficiency in previous studies.

**Motor strength, gait and co-ordination**

##### Grip Strength

Grip strength of each animal was measured with a grip strength meter (GT3, Griptest v3.48, Bioseb). The animal was placed onto a grid and gently pulled backward by the tail to measure peak resistance force. The average force over six trials was recorded and normalised according to their body weight (g).

##### **Rotarod**

The animals were first trained on the rotarod (Panlab, Harvard Apparatus) at a constant speed of 4rpm, until they were able to stay on for 60 seconds. Latency to fall was tested over 3 trials, with the rod accelerating from 4 to 40 rpm over 120 seconds, and 1-minute break in between each test.

##### **Beam Walk Balance Test**

Firstly, mice were trained in walking across two elevated narrow beams – one 12 mm wide and one 6 mm wide (Balance Beam, Maze Engineers) - connected to two platforms either end. Mice crossed each beam three times as part of the training. Next, the test was conducted and the time taken for mice to cross each beam between two marked points was measured. All mice crossed each beam twice and the average time was calculated. Any paw slips or hindlimb dragging were recorded too and, together with the crossing times, were used to score the motor performance of each mouse.^91^ Mice received a score between 1-7 on their beam-walking ability. Score = 1 indicates (a complete inability to cross the beam) and 7 (= best walking ability). An intermediate score reflected an ability to cross the beam but with varying degrees of slips and hindlimb dragging.

##### **CatWalk XT Gait analysis**

Gait analysis was performed on WT, *mdx52*, and treated *mdx52.µDys* at 12 weeks of age by using the automated gait analysis CatWalk XT system (Noldus, Wageningen, The Netherlands). Each mouse was individually placed at one end of the CatWalk and allowed to freely traverse a backlit, filmed section of the walkway. A minimum of three successful runs were recorded per session, with a run considered successful if it met the standard criteria. Paw prints from these successful runs were captured, classified, and analyzed using CatWalk XT software v10.7 (Noldus) to generate comprehensive gait measurements. Parameters recorded included the stride length (the distance between successive paw placement of the same paw in cm), swing speed (speed of the paw between successive paw placement, cm/sec), duration of the run (seconds), and regularity index (% index for the degree of interlimb coordination during gait).^92^

The mean print position is the distance between the position of the hind paw and the position of the previously placed front paw on the ipsilateral side and in the same step cycle. A positive value of the print position indicates that the hind paw is placed behind the front paw. A negative value of the print position indicates that the hind paw is placed in front of the front paw as sign of incoordination.

### **Tissue collection**

The gene therapy study mice were euthanised by CO_2_ at 12 weeks old. The right brain hemisphere was microdissected to cortex, hippocampus, midbrain, hindbrain and cerebellum for protein and molecular analysis. The left hemisphere was rinsed with PBS before OCT embedding in dry ice bath with 100% ethanol. Sample were stored at -70^o^C until further analysis. Peripheral tissues heart, liver, tibialis anterior were harvested for protein, molecular and immunofluorescence analysis.

For GFP gene biodistribution studies, mice were perfused under isoflurane anaesthesia and GFP native fluorescence imaged by stereoscopic microscopy.^45^ Brains were fixed in 4%PFA and sectioned at 40 microns for free floating whole brain immunostaining.^45^

### **Brain Immunofluorescence**

For *mdx52* fresh frozen sagittal brain were sectioned Leica CM 1860 UV cryostat at 20 µm (n=5) and were stored at -70^o^C. For immunofluorescence sections were thawed at room temperature for 30 seconds before incubation in100% methanol for 2 minutes at -20^o^C. They were then washed with PBS and incubated in 2% goat serum, 0.3% triton X-100, and 1% bovine serum albumin diluted in PBS for 40 minutes at room temperature. They were then incubated overnight at 4^o^C with primary antibodies (**Supplementary Table 1)**. Sections were washed with PBS and incubated for 1 hour at room temperature with secondary antibodies (**Supplementary Table 1**). The images were taken on Zeiss Observer 7 colour fluorescent microscope. Free floating GFP brain intracranial biodistribution immunofluorescence undertaken with using Ab Cam GFP 1:10000 as previously published.^45^

### **Muscle Immunofluorescence**

Frozen muscle sections were stained with MANEX 1B (Glenn Morris) followed by a secondary antibody, Rabbit Anti-IgG1 + IgG2a + IgG3 (Ab133469 Abcam). a-Bungarotoxin (B13422 Thermofisher) was then applied before nuclear counterstain with Hoechst 33342 (H1399 ThermoFisher). Detailed description of the protocol is described in supplementary material.

### **Simple Capillary Western Blot (WES)**

Protein samples were extracted from 12-week-old mice as described previously.^18^￼ Simple capillary western blot (WES, Bio-techne) was used to quantify dystrophin proteins in brain muscle and heart. A total of 2 μg protein was loaded for 66-440 kDa separation and the total protein detection (Bio-techne). The antibodies used are described in **Supplementary Table 1**. μDys and dystrophin interactors levels were normalised to the total protein.

### **μDys mRNA transcript and Vector genomic copies quantification**

Samples were lysed with Trizol (ThermoFisher) and Tungsten beads (Qiagen) using the QIAGEN Tissue Lyser II and extracted with the PureLink RNA Mini Kit (Invitrogen). RNA was normalised and cDNA was generated with the High-Capacity cDNA Reverse Transcription kit (Applied Biosystems). Primers and probe (forward: 5’-GTGTTCAGGTCTTCCAGAGTG, reverse: 5-GGAGAAATTGCGCCTCTGA, probe: 5’- /56FAM/TTT GGG CAT/ZEN/TCA GCT CTC ACC GTA/3IABkFQ/ -3’) were designed to target Exon 60 present in μDys transgene with mouse GAPDH as housekeeping gene. DNA was extracted from tissue with DNeasy Blood & Tissue Kit (QIAGEN) to quantify vector genome copies (vgc) by TaqMan Multiplexed qPCR with Luna Universal Probe qPCR Master Mix on QuantStudio 5.

### **Microscopy**

### High-resolution images of the stained tissues were acquired using the NanoZoomer S20 scanner (Hamamatsu) at 40× magnification. Image analysis was performed with NDP software (Hamamatsu). GFP images were taken on Leica DM 4000.

Supplementary Table 1. **Antibodies list**

| **Antibody** | **Species** | **Dilution** | **Primary/Secondary** | **Company** |
| --- | --- | --- | --- | --- |
| NCL-DYS3 | Mouse | 1:20 (IHC) | Primary | Leica Biosystems, Cat. No. NCL-DYS3 |
| β-dystroglycan | Rabbit | 1:200 (IHC) | Primary | Proteintech, Cat No. 11017-1-AP |
| Syntrophin beta 1 | Rabbit | 1:500 (IHC) | Primary | Abcam, ab242046 |
| GABA_A_R | Rabbit | 1:400 (IHC) | Primary | Proteintech Cat No. 14104-1-AP |
| Gephyrin | Rabbit | 1:400 (IHC) | Primary | Proteintech, 12681-1-AP |
| VGLUT1 | Rabbit | 1:200 (IHC) | Primary | Proteintech |
| MANEX1 | Mouse | 1:25 (IHC) | Primary | Glenn Morris |
| Dystrophin | Mouse | 1:1500 (WES) | Primary | Abcam, Cat No. Ab154168 |
| β-dystroglycan | Mouse | 1:200 (WES) | Primary | Leica, Cat No. NCL-b-Dg |
| Syntrophin beta 1 | Rabbit | 1:200 (WES) | Primary | Abcam, Cat No. ab242046 |
| GABA_A_R | Rabbit | 1:200 (WES) | Primary | Proteintech, Cat No. 14104-1-AP |
| Gephyrin | Rabbit | 1:400 (WES) | Primary | Proteintech, 12681-1-AP |
| VGLUT1 | Rabbit | 1:1500 (WES) | Primary | Proteitech, Cat No. 29469-1-AP |
| anti-mouse IgG2A Alexa fluorophore 594 | Goat | 1:1000 (IHC) | Secondary | Invitrogen, A-211350 |
| anti-rabbit Alexa fluorophore 488 | Goat | 1:1000 (IHC) | Secondary | Invitrogen, A-11008 |

**Supplementary Figures**

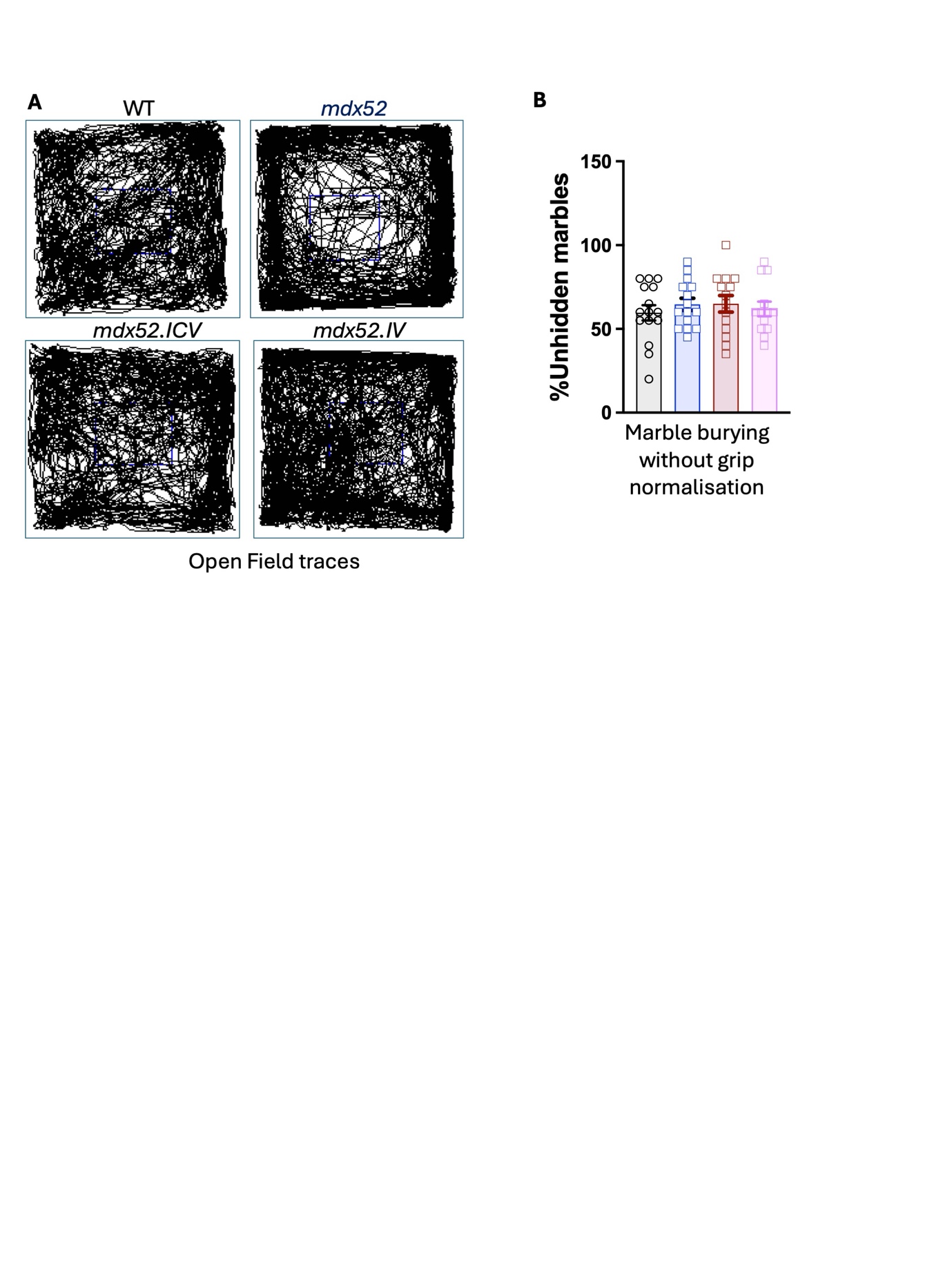
**Supplementary Figure 1 Gene therapy effects on behavioural parameters (A)** Open field representative traces of motor activity showing increased periphery traces in control mdx52 compared to WT and treated *mdx52* demonstrating improvement in thigomataxis behavioura **(B)** Percentage of unhidden marbles in marble test. Results presented as mean ± SEM analysed with one-way ANOVA with Tukey’s post-hoc.

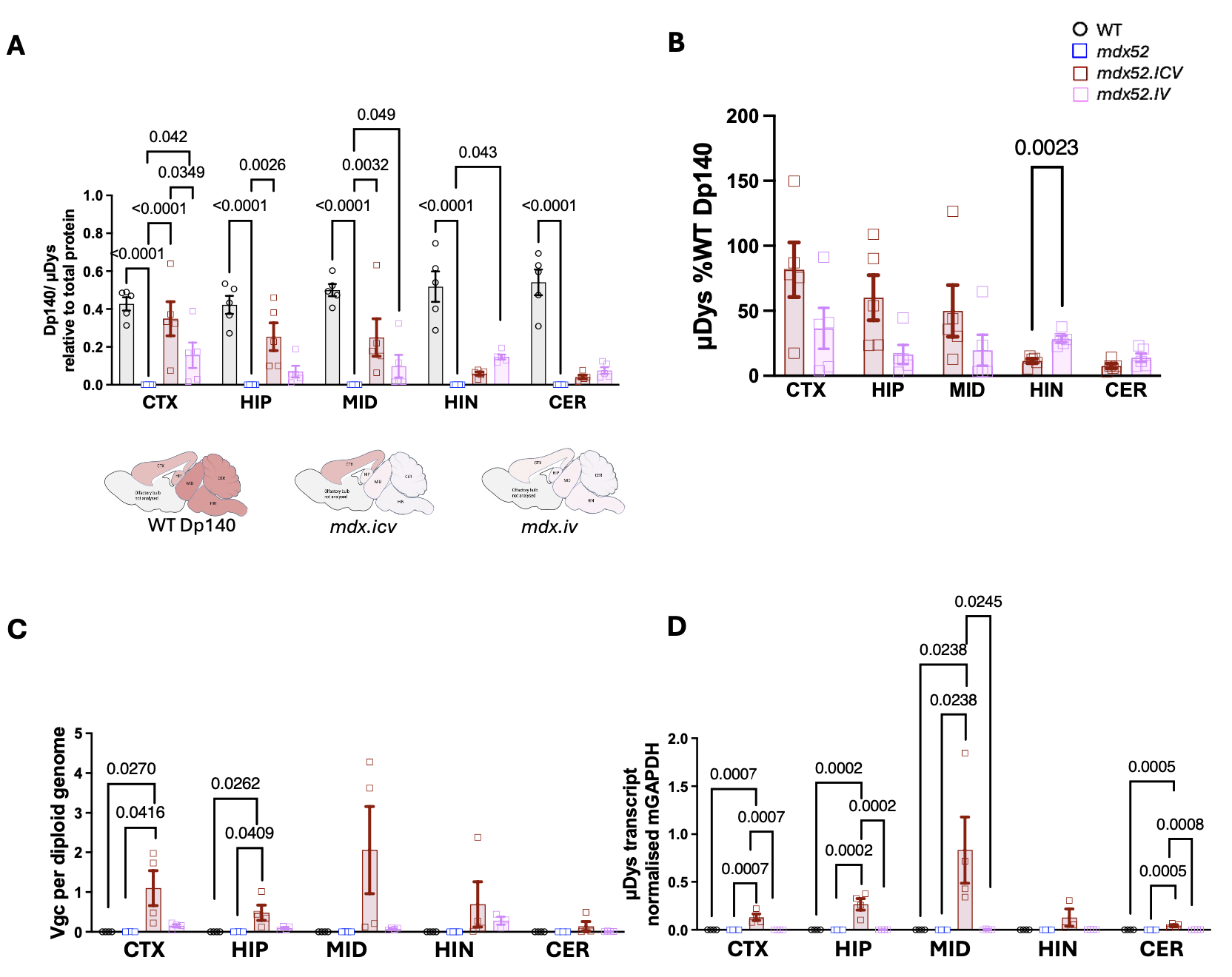

**Supplementary Figure 2 Neonatal ICV and IV AAV9.μDys gene therapy expression.** Brains collected 12 weeks post-gene therapy and dissected into cortex, hippocampus, midbrain, hindbrain and cerebellum **(A)** μDys protein concentration relative to WT mice assessed by simple western capillary (n=5). WT represents Dp140 levels **(B)** μDys levels as %WTDp140 **(C)** Vector genome copies quantified by Taqman Multiplex qPCR (n=4) **(D)** Micro-dystrophin transcript expression relative to mouse GAPDH assessed by RT TaqMan Multiplex qPCR (n=4). Adjusted p-value for significant groups indicated in graphs, analysed by mean comparison to *mdx52.ICV* and *mdx52.IV* with a one-way ANOVA, followed by Dunnet’s test (p-value<0.05) CTX: cortex, HIP, hippocampus, MID: midbrain, HIN: hindbrain, CER: cerebellum

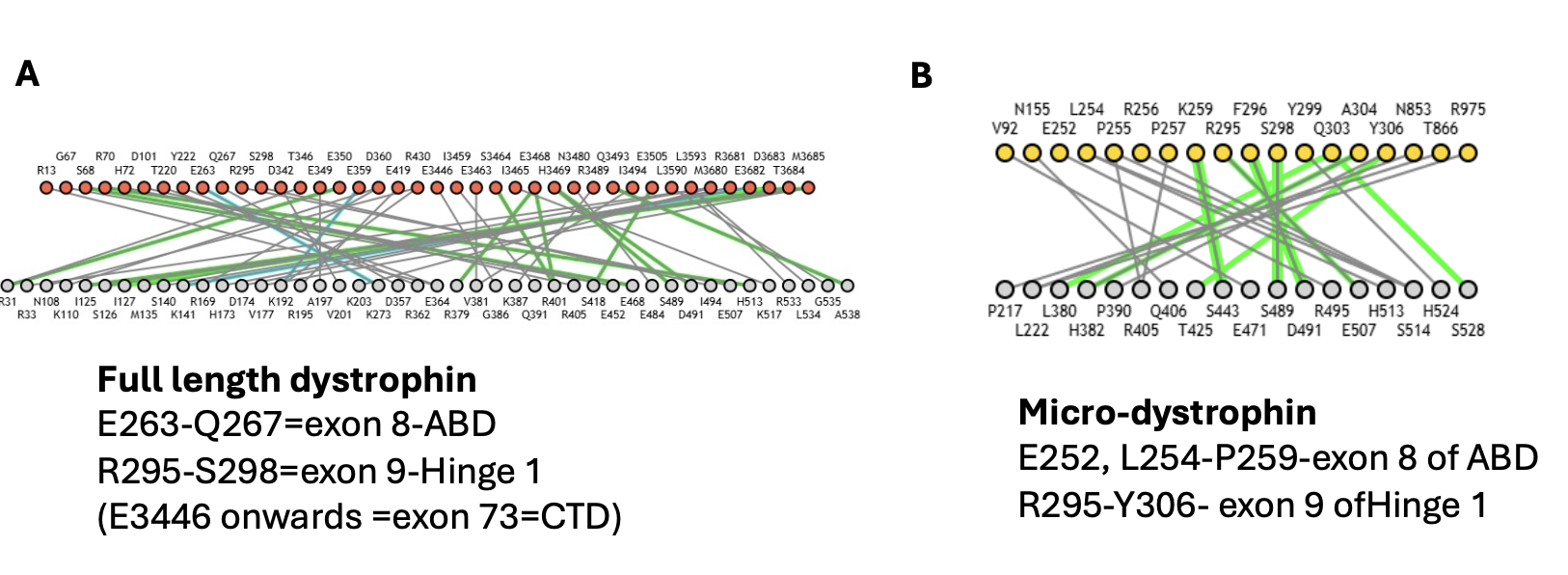

**Supplementary Figure 3 Schematic SNTB1 interaction with Dp427 µDys with common amino acid positions listed (A)** Interaction of full-length dystrophin **(B)** Micro-dystrophin domains of interactions

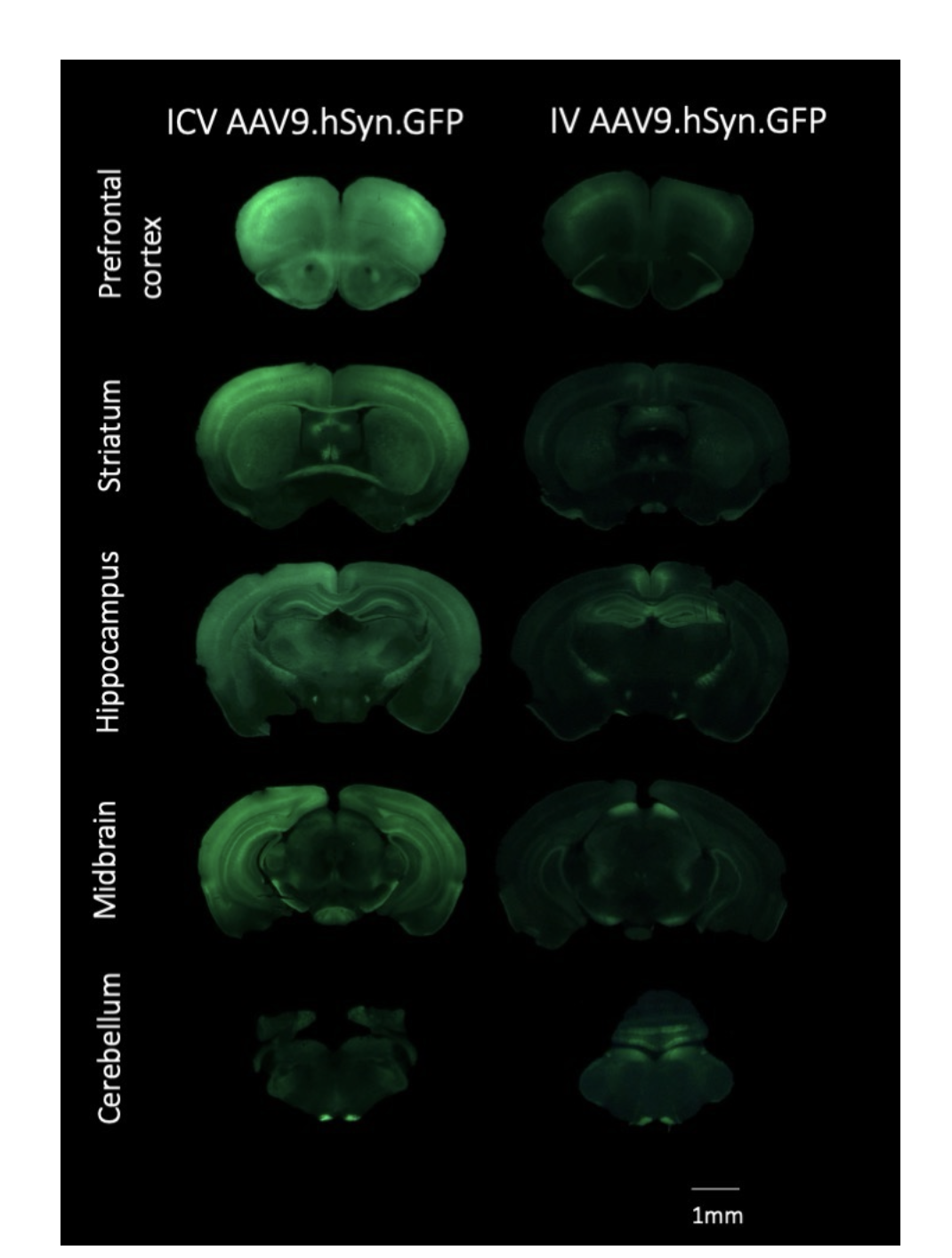
**Supplementary Figure 4 AAV9.hSyn.GFP brain transduction.** Brain GFP immunofluorescence and stereoscopic fluorescence images after vector delivered to P1 wild-type pups ICV or IV. Inserts: 1mm

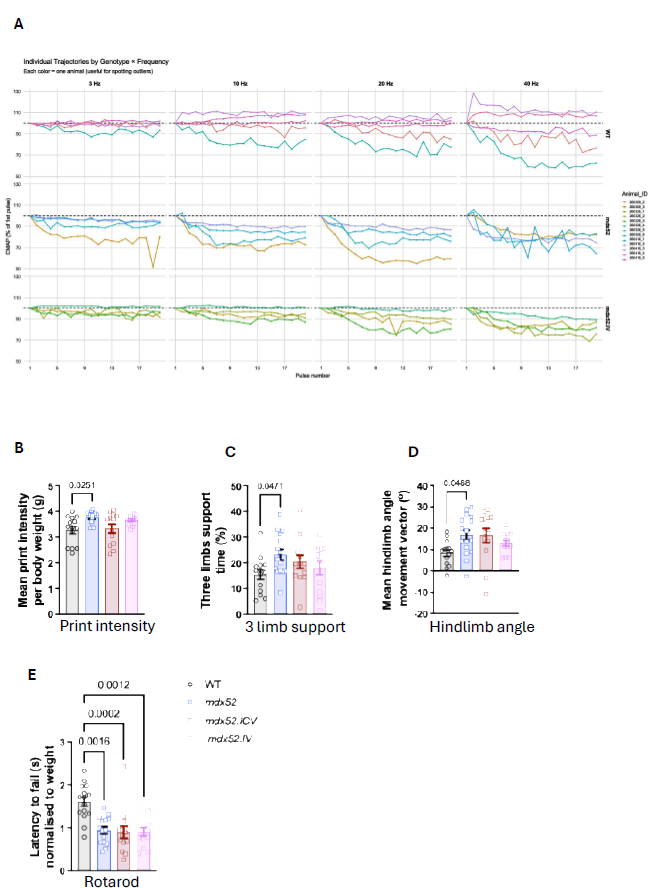

**Supplementary Figure 5 AAV9.μDys gene therapy improves motor coordination (A)** RNS at 3, 10, 20, 40Hz representing the individual animals used. **(B, C)** Mean print position, mean print intensity normalised to body weight and max print intensity. **(D)** Quantification of percentage of three limb support time **(E)** Mean hindlimb angle movement vector **(J)** Average time spent on the rotarod in seconds normalised to body weight analysed by one-way ANOVA with Tukey’s post-hoc. Results presented as mean ± SEM.

**Supplementary Table 2. CatwalkXT parameters**

| **Category** | **Parameter** | **WT** | **mdx52** | **mdx52 ICV** | **mdx52 IV** |
| --- | --- | --- | --- | --- | --- |
| Run characteristics | Mean speed (cm/s) | 21.39 ± 1.51 | 15.22 ± 1.6 | 18.6 ± 2.39 | 17.69 ± 1.96 |
|  | Mean maximum speed variation (%) | 41.21 ± 4.15 | 58.49 ± 4.64 | 49.45 ± 4.95 | 49.18 ± 5.36 |
|  | Cadence | 14. ± 0.68 | 12.42 ± 0.75 | 13.12 ± 0.91 | 12.52 ± 0.75 |
|  | Number of steps | 73.36 ± 3.56 | 73.33 ± 3.88 | 72.77 ± 3.33 | 77.15 ± 3.73 |
| Temporal | Step cycle (s) | 0.29 ± 0.02 | 0.33 ± 0.02 | 0.32 ± 0.02 | 0.32 ± 0.02 |
|  | Stand (s) - all limbs | 0.16 ± 0.01 | 0.22 ± 0.02 | 0.19 ± 0.02 | 0.20 ± 0.02 |
|  | Swing (s) - all limbs | 0.13 ± 0.01 | 0.13 ± 0.01 | 0.13 ± 0.01 | 0.13 ± 0.01 |
|  | Swing speed (cm/s) | 51.74 ± 2.4 | 46.48 ± 2.56 | 49.06 ± 3.68 | 47.15 ± 3.03 |
|  | Duty cycle (%) - all limbs | 53.90 ± 0.94 | 57.03 ± 0.86 | 56.61 ± 1.32 | 55.88 ± 1.57 |
|  | Initial dual stance (s) | 0.02 ± 0.01 | 0.04 ± 0.01 | 0.03 ± 0.01 | 0.03 ± 0.01 |
|  | Terminal dual stance (s) | 0.02 ± 0.01 | 0.04 ± 0.01 | 0.02 ± 0.01 | 0.04 ± 0.01 |
| Spatial | Stride length (cm) | 6.18 ± 0.16 | 5.75 ± 0.17 | 5.83 ± 0.26 | 5.77 ± 0.22 |
|  | Base of support (mm) - forelimbs | 12.73 ± 0.28 | 13.99 ± 0.40 | 13.95 ± 0.48 | 13.64 ± 0.37 |
|  | Print position (mm) | 5.18 ± 0.84 | 9.00 ± 1.23 * | 6.15 ± 0.75 | 6.86 ± 0.68 |
|  | Mean hindlimb  angle body axis (o) | 5.07 ± 1.69 | 13.27 ± 1.96 * | 13.69 ± 2.64 ** | 11.75 ± 1.75 |
|  | Mean hindlimb  angle movement vector (o) | 8.4 ± 1.56 | 16.77 ± 2.33 * | 16.55 ± 3.35 | 13. ± 1.36 |
| Postural support | Single support time (%) | 4.84 ± 1.06 | 2.94 ± 0.58 | 3.32 ± 0.74 | 4.49 ± 1.10 |
|  | Diagonal support time (%) | 70.60 ± 2.22 | 63.00 ± 2.74 | 66.21 ± 3.28 | 66.74 ± 3.03 |
|  | Girdle support time (%) | 1.52 ± 0.30 | 1.83 ± 0.24 | 2.19 ± 0.28 | 1.77 ± 0.32 |
|  | Lateral support time (%) | 2.45 ± 0.49 | 1.49 ± 0.29 | 1.04 ± 0.32 | 1.54 ± 0.37 |
|  | Three limbs support time (%) | 15.44 ± 1.9 | 23.22 ± 2.13 * | 20.42 ± 2.52 | 18.12 ± 2.69 |
|  | Four limbs support time (%) | 5.02 ± 1.18 | 7.52 ± 1.54 | 6.58 ± 1.36 | 7.28 ± 1.94 |
| Inter-limb co-ordination | Regularity Index (%) | 93.96 ± 1.34 | 95.23 ± 0.78 | 95.32 ± 1.33 | 94.33 ± 1.21 |
|  | Number of step sequence | 15.14 ± 0.74 | 15.60 ± 0.86 | 15.46 ± 0.69 | 16.08 ± 0.78 |
|  | Step sequence (%) CA | 21.02 ± 3.05 | 26.73 ± 4.66 | 21.41 ± 4.51 | 23.34 ± 4.74 |
|  | Step sequence (%) CB | 21.08 ± 3.25 | 21.94 ± 4.14 | 23.55 ± 5.55 | 17.75 ± 2.41 |
|  | Step sequence (%) AA | 24.5 ± 4.81 | 12.61 ± 3.39 | 12.9 ± 3.44 | 19.52 ± 4.36 |
|  | Step sequence (%) AB | 30.06 ± 6.17 | 37.82 ± 4.72 | 41.23 ± 5.91 | 38.98 ± 6.36 |
|  | Step sequence (%) RA | 2.42 ± 1.36 | 0.51 ± 0.51 | 0.91 ± 0.62 | 0.00 ± 0.00 |
|  | Step sequence (%) RB | 0.92 ± 0.63 | 0.39 ± 0.39 | 0.00 ± 0.00 | 0.40 ± 0.20 |
|  | Coupling (%) LF>RH | 57.36 ± 12.96 | 21.41 ± 10.05 | 24.8 ± 11.53 | 25.2 ± 11.48 |
|  | Coupling (%) RH>LF | 56.27 ± 12.85 | 77.55 ± 10.04 | 74.95 ± 11.54 | 74.36 ± 11.42 |
|  | Coupling (%) RF>LH | 42.2 ± 13.03 | 47.63 ± 12.19 | 25.37 ± 11.41 | 46.67 ± 13.22 |
|  | Coupling (%) LH>RF | 49.94 ± 13. | 58.28 ± 12.17 | 74.03 ± 11.35 | 52.31 ± 13.2 |
|  | Coupling (%) LF>LH | 49.05 ± 0.76 | 51.64 ± 1.23 | 52.6 ± 0.80 | 51.36 ± 1.28 |
|  | Coupling (%) LH>LF | 51.6 ± 0.88 | 48.43 ± 1.22 | 47.13 ± 0.81 | 48.43 ± 1.16 |
|  | Coupling (%) RF>RH | 50.94 ± 1.09 | 50.91 ± 0.95 | 52.31 ± 0.69 | 51.47 ± 1.18 |
|  | Coupling (%) RH>RF | 50.05 ± 1.11 | 49.88 ± 1.19 | 48.33 ± 0.75 | 48.38 ± 1.23 |
|  | Coupling (%) LF>RF | 50.04 ± 0.51 | 50.7 ± 1.01 | 50.19 ± 0.93 | 50.44 ± 0.96 |
|  | Coupling (%) RF>LF | 50.23 ± 0.59 | 48.88 ± 1. | 50.12 ± 0.82 | 48.88 ± 0.91 |
|  | Coupling (%) LH>RH | 50.96 ± 1.16 | 49.93 ± 1.02 | 48.54 ± 0.96 | 50.65 ± 0.68 |
|  | Coupling (%) RH>LH | 48.99 ± 0.77 | 49.8 ± 0.96 | 51.29 ± 1.07 | 48.62 ± 0.67 |
| Paw loading | Mean print intensity per body weight (g) - all limbs | 3.28 ± 0.15 | 3.75 ± 0.06 * | 3.34 ± 0.17 | 3.66 ± 0.05 |
|  | Mean print intensity per body weight (g) - hindlimbs | 3.55 ± 0.18 | 3.98 ± 0.07 | 3.52 ± 0.20 | 3.89 ± 0.06 |
|  | Mean print intensity per body weight (g) - forelimbs | 3.01 ± 0.12 | 3.54 ± 0.06 ** | 3.15 ± 0.15 † | 3.42 ± 0.06 * |
|  | Maximum print intensity per body weight (g) - all limbs | 5.73 ± 0.13 | 6.41 ± 0.09 * | 6.04 ± 0.17 † | 6.11 ± 0.05 |
|  | Maximum print intensity per body weight (g) - hindlimbs | 3.40 ± 0.18 | 3.80 ± 0.07 | 3.40 ± 0.20 | 3.70 ± 0.06 |
|  | Maximum print intensity per body weight (g) - forelimbs | 2.90 ± 0.13 | 3.40 ± 0.05 * | 3.00 ± 0.16 † | 3.30 ± 0.06 * |
| Paw contact | Print area per body weight (cm2/g) - all limbs | 0.009 ± 0.001 | 0.011 ± 0.001 | 0.013 ± 0.001 * | 0.010 ± 0.001 |
|  | Print area per body weight (cm2/g)- hindlimbs | 0.009 ± 0.001 | 0.011 ± 0.001 | 0.012 ± 0.001 * | 0.009 ± 0.001 § |
|  | Print area per body weight (cm2/g)- forelimbs | 0.010 ± 0.001 | 0.012 ± 0.001 | 0.013 ± 0.001 | 0.011 ± 0.001 |
|  | Mean print width per body weight (cm/g) | 0.024 ± 0.001 | 0.026 ± 0.001 * | 0.026 ± 0.001 | 0.025 ± 0.001 |
|  | Mean hindlimb width per body weight (cm/g) | 0.022 ± 0.001 | 0.026 ± 0.001 * | 0.026 ± 0.001 * | 0.023 ± 0.001 |
|  | Mean forelimb width per body weight (cm/g) | 0.026 ± 0.001 | 0.027 ± 0.001 | 0.026 ± 0.000 | 0.026 ± 0.000 |
|  | Mean print length per body weight (cm/g) | 0.028 ± 0.001 | 0.030 ± 0.001 | 0.031 ± 0.001 | 0.028 ± 0.001 |
|  | Mean hindlimb length per body weight (cm/g) | 0.027 ± 0.001 | 0.029 ± 0.001 | 0.031 ± 0.001 | 0.028 ± 0.001 |
|  | Mean forelimb length per body weight (cm/g) | 0.028 ± 0.001 | 0.030 ± 0.001 | 0.030 ± 0.001 | 0.029 ± 0.000 |
|  | Maximum contact area per body weight (cm2/g) - all limbs | 0.008 ± 0.001 | 0.009 ± 0.001 | 0.010 ± 0.001 * | 0.008 ± 0.001 § |
|  | Maximum contact area per body weight (cm2/g) - hindlimbs | 0.007 ± 0.001 | 0.009 ± 0.001 | 0.010 ± 0.001 | 0.007 ± 0.001 |
|  | Maximum contact area per body weight (cm2/g) - forelimbs | 0.008 ± 0.001 | 0.009 ± 0.001 | 0.011 ± 0.001 | 0.008 ± 0.001 |
|  | Mean hindlimb toe spread (mm) | 7.43 ± 0.22 | 8.56 ± 0.16 *** | 8.44 ± 0.21 ** | 8.07 ± 0.20 |
|  | Hindlimb toe spread (mm) - right | 7.63 ± 0.27 | 8.10 ± 0.22 | 8.50 ± 0.19 * | 7.98 ± 0.20 |
|  | Hindlimb toe spread (mm) - left | 7.33 ± 0.24 | 8.87 ± 0.18 **** | 8.39 ± 0.28 * | 8.12 ± 0.25 |
|  | Mean hindlimb internal toe spread (mm) | 4.43 ± 0.20 | 5.11 ± 0.13 * | 5.15 ± 0.09 ** | 4.71 ± 0.13 |
|  | Hindlimb internal toe spread (mm) - right | 4.41 ± 0.18 | 5.00 ± 0.23 | 5.30 ± 0.14 †† | 4.55 ± 0.12 |
|  | Hindlimb internal toe spread (mm) - left | 4.42 ± 0.28 | 5.21 ± 0.14 * | 5.02 ± 0.14 | 4.87 ± 0.19 |

Data presented as mean ± SEM and analysed with one-way ANOVA with Tukey’s post-hoc.

**mdx52* untreated mice worse gait function than WT, or improvement after treatment

** *mdx52* ICV treated mice worse gait function than WT, or IV improved compared to ICV

Supplementary Video legends:

SV1: Control WT - fluid gait and direct trajectory

SV2: Control *mdx52* -slower gait, brief pauses and weaving trajectory.

SV3: *mdx.icv* gait - improvement in fluidity and direct trajectory, no waddle.

SV4: *mdx.iv* gait- improvement in fluidity, speed and direct trajectory with mild waddle.
